# Target DNA-dependent activation mechanism of the prokaryotic immune system SPARTA

**DOI:** 10.1101/2023.09.23.559129

**Authors:** Giada Finocchio, Balwina Koopal, Ana Potocnik, Clint Heijstek, Martin Jinek, Daan C. Swarts

**Author notes:** Equal contribution (Giada Finocchio, Balwina Koopal).

## Abstract

In both prokaryotic and eukaryotic innate immune systems, TIR domains function as NADases that degrade the key metabolite NAD^+^ or generate signaling molecules. Catalytic activation of TIR domains requires oligomerization, but how this is achieved varies in distinct immune systems. In the <u>S</u>hort <u>p</u>rokaryotic <u>Ar</u>gonaute (pAgo)/<u>T</u>IR-<u>A</u>PAZ (SPARTA) immune system, TIR NADase activity is triggered upon guide RNA-mediated recognition of invading DNA by an unknown mechanism. Here, we describe cryo-EM structures of SPARTA in the inactive monomeric and target DNA-activated tetrameric states. The monomeric SPARTA structure reveals that in the absence of target DNA, a C-terminal tail of TIR-APAZ occupies the nucleic acid binding cleft formed by the pAgo and TIR-APAZ subunits, suppressing SPARTA activation. In the active tetrameric SPARTA complex, guide RNA-mediated target DNA binding displaces the C-terminal tail and induces conformational changes in pAgo that facilitate SPARTA-SPARTA dimerization. Concurrent release and rotation of one TIR domain allow it to form a composite NADase catalytic site with the other TIR domain within the dimer, and generate a self-complementary interface that mediates cooperative tetramerization. Combined, this study provides critical insights into the structural architecture of SPARTA and the molecular mechanism underlying target DNA-dependent oligomerization and catalytic activation.

**KEY POINTS:**

- Inactive monomeric SPARTA is autoinhibited by the C-terminal tail of TIR-APAZ
- Target DNA recognition causes C-terminal tail release, pAgo restructuring, and dimerization
- TIR domain rotation enables catalytic activation and cooperative tetramer formation

## INTRODUCTION

Argonaute proteins comprise a diverse family of proteins that confer immunity in all domains of life^1, 2^. Eukaryotic Argonautes (eAgos) associate with short (15-30 nt) RNA guides to target complementary RNA sequences. Catalytically active ‘slicing’ eAgos can cleave the targeted RNA, for example during antiviral defense, while catalytically inactive eAgos recruit accessory proteins to the targeted RNA to elicit transcriptional or post-transcriptional gene silencing^3^. As such, eAgos can regulate native gene expression but also interfere with virus and transposon propagation^1^. In contrast, archaea and bacteria encode a large variety of prokaryotic Argonaute proteins (pAgos) that, besides reported involvement in DNA replication^4, 5^ and DNA repair^6^, mainly act as prokaryotic immune systems that interfere with invading plasmid and bacteriophage DNA^7–13^.

Based on their phylogeny, pAgos are subdivided in long-A, long-B, and short pAgos^14, 15^. eAgos and long pAgos share a bilobed structural architecture^16^. The MID-PIWI lobe consists of the middle (MID) and p-element induced wimpy testis (PIWI) domains^17^, and is responsible for guide 5′-end binding and guide-mediated target binding^17–20^. This lobe is connected through the linker 2 (L2) motif to the N-terminal lobe, which is comprised of the N-terminal (N) domain, the linker 1 (L1) motif and the PAZ (PIWI-ARGONAUTE-ZWILLE) domain. These domains have been implicated in guide loading, guide 3’ end binding, target binding specificity, and Ago protein turnover^18, 20–22^. In contrast to eAgos and long pAgos, short pAgos are comprised of the MID and PIWI domains only, and all short pAgos are catalytically inactive^14^. Short pAgos are encoded in operons together with (and sometimes fused to) proteins comprising an Analog of PAZ (APAZ) domain, which displays homology to the N domain of long Agos^15^. APAZ domains are fused to one of various ‘effector’ domains including (but not limited to) Sirtuin (Sir2), RecG/DHS-like, Mrr-like, DUF4365, and Toll/interleukin-1 receptor (TIR) domains^14, 15^. Upon guide RNA-mediated recognition of invading plasmid or bacteriophage DNA, short pAgo/TIR-APAZ (SPARTA), short pAgo/Sir2-APAZ (SPARSA), and short pAgo/DUF4365 (SPARDA) systems trigger abortive infection^11–13^. SPARTA and SPARSA systems achieve abortive infection through enzymatic degradation of the central metabolite nicotinamide adenine dinucleotide (NAD^+^)^12, 13^.

The SPARTA TIR domain is homologous to other TIR effector domains found in prokaryotic, plant, and animal immune systems^12, 23–33^. In these immune systems, TIR domains function in conjunction with variable ‘sensor’ proteins that assemble into higher-order structures upon recognition of a pathogen-derived molecular signature, thereby facilitating TIR domain oligomerization^23, 25, 28, 31, 33, 34^. Not all TIR domains have an enzymatic function; in scaffolding TIR domains, TIR oligomerization results in formation of signalosomes^35^. In enzymatic TIR NADases, TIR oligomerization triggers catalytic activation whereby head-to-tail TIR domain interactions allow one TIR domain to insert its flexible ‘BB-loop’ into a pocket of a second TIR domain. This forms a composite catalytic site capable of NAD^+^ binding and hydrolysis^25, 28, 31, 33, 34^. Prokaryotic TIR effectors degrade NAD^+^ either to generate secondary messenger molecules^26^ or to trigger cell death and consequentially prevent invader propagation^12, 27–30^. The structural basis for TIR domain oligomerization has been described for various eukaryotic and prokaryotic immune systems^23, 25, 28, 31, 34^. Notably, the interaction interfaces underlying TIR oligomerization are variable for distinct immune systems, which gives rise to various TIR oligomerization modes. In SPARTA systems, RNA-mediated target DNA binding results in tetramerization of the heterodimeric pAgo:TIR-APAZ complex, suggesting that the TIR domains are activated upon oligomerization^12^. However, the molecular mechanisms underlying SPARTA activation remain unknown.

Here, we report cryo-EM structures of the *Bacillales bacterium* (Bab) SPARTA system in its inactive monomeric and activated tetrameric forms. The structure of the monomeric BabSPARTA complex reveals an autoinhibited state in which nucleic acid binding channel formed by the interaction of pAgo and TIR-APAZ subunits is occupied by the C-terminal tail (Ct) of TIR-APAZ. Interactions of the Ct and the MID domain with TIR-APAZ keep TIR in an inhibited conformation. The structure of the activated BabSPARTA tetramer reveals that RNA-guided target DNA binding displaces the Ct and induces conformational changes that result in pAgo-mediated dimerization. This in turn releases one TIR domain and enables dimerization with the TIR domain of the other protomer, generating a composite NADase catalytic site. The TIR-TIR dimer is further stabilized through cooperative interactions with the TIR-TIR dimer of another activated SPARTA dimer, resulting in the formation of a catalytically active tetramer. Together, these insights reveal the mechanisms underlying the RNA-guided target DNA recognition and catalytic activation of SPARTA systems.

## RESULTS

### Structural basis for short pAgo-APAZ heterodimerization

To gain insights into the molecular mechanism of the SPARTA system, we reconstituted the BabSPARTA complex (**Figure 1A**) together with a 21-nucleotide guide RNA in the absence of a target DNA for structural analysis by cryogenic electron microscopy. The resulting reconstruction, determined at a resolution of 2.6 Å, reveals a monomeric, binary BabAgo:BabTIR-APAZ SPARTA complex (**Figure 1B, C**, **Supplementary Figure S1**, and **Table S1**). This monomeric SPARTA complex reconstruction lacks interpretable density for the guide RNA (**Supplementary Figure S1D**), suggesting that inactive SPARTA binds guide RNAs with a low affinity or that SPARTA-bound guide RNAs are conformationally heterogeneous in the absence of target DNA. Alternatively, SPARTA systems might bind preformed guide RNA-target DNA duplexes or rely on other mechanisms for efficient guide RNA loading.

**Figure 1.**
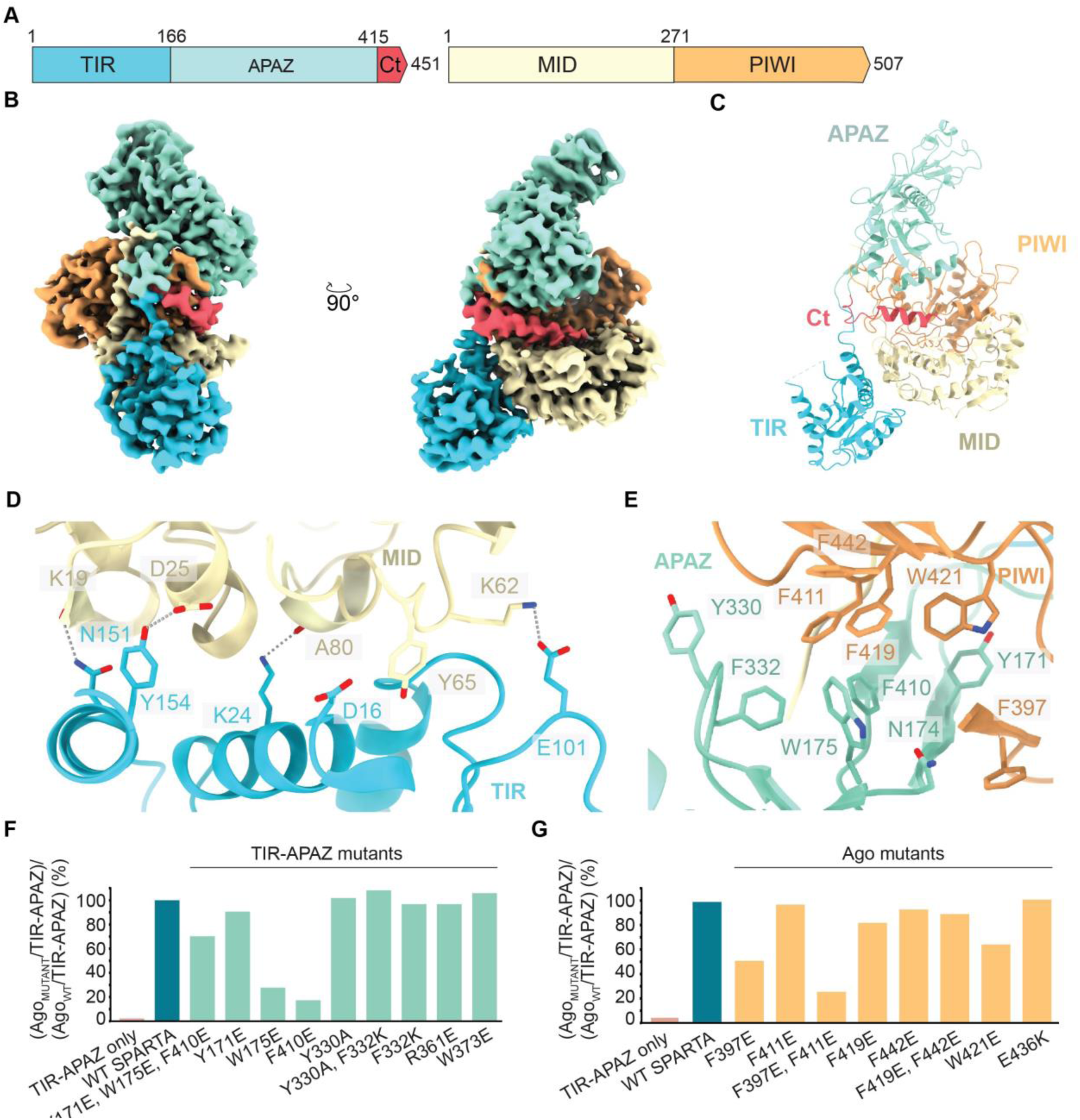
Molecular architecture of the monomeric apo-SPARTA complex. (**A**) Schematic diagram of the domain organization of the BabTIR-APAZ and BabAgo proteins. TIR, Toll-interleukin-1 receptor-like; APAZ, analog of PIWI-AGO-ZWILLE domain; Ct, C-terminal tail; MID, Middle; PIWI, P-element induces wimpy testis domain. (**B**) Cryo-electron microscopic (cryo-EM) densities of the BabSPARTA complex. Domains are colored according to the scheme in (A). (**C**) Cartoon representation of the overall structure of the Apo-BabSPARTA complex. Domains are colored according to the scheme in (A). (**D**) Close-up view of MID-TIR interactions. (**E**) Close-up view of PIWI-APAZ interactions. (**F and G**) Multiple residues at the PIWI-APAZ interface contribute to heterodimerization of TIR-APAZ and pAgo. 6xHis-MBP-BabTIR-APAZ or mutants thereof (F) were co-expressed in *E. coli* with BabAgo or mutants thereof (G) and proteins were purified using amylose affinity chromatography. Graphs show the percentage of the BabAgo/BabTIR-APAZ ratio normalized against the BabAgo/BabTIR-APAZ ratio of the WT proteins.

The monomeric BabSPARTA structure reveals that the BabTIR-APAZ subunit wraps around the BabAgo subunit (**Figure 1B, C**). BabAgo adopts a compact fold with a canonical arrangement of MID and PIWI domains, as previously observed in other pAgo and eAgo proteins^19, 36–38^ (**Supplementary Figure S2**). BabTIR-APAZ adopts an extended conformation, with the TIR and APAZ domains contacting the BabAgo MID and PIWI domains, respectively (**Figure 1C**). Structural superposition of the BabSPARTA complex with the long-A pAgo from *Thermus thermophilus* (TtAgo) reveals that the APAZ domain resembles the N-domain of long-A pAgo proteins (**Supplementary Figure S2**). The TIR-MID domain interface extends over 633 Å^2^ and features a network of salt-bridge and hydrogen-bonding interactions (**Figure 1D**). The TIR-MID interaction sterically precludes the TIR domain from forming head-to-tail TIR-TIR interactions that are typically required for their catalytic activation ^23, 28, 31, 34^, explaining why apo-SPARTA is catalytically inactive.

**Figure 2.**
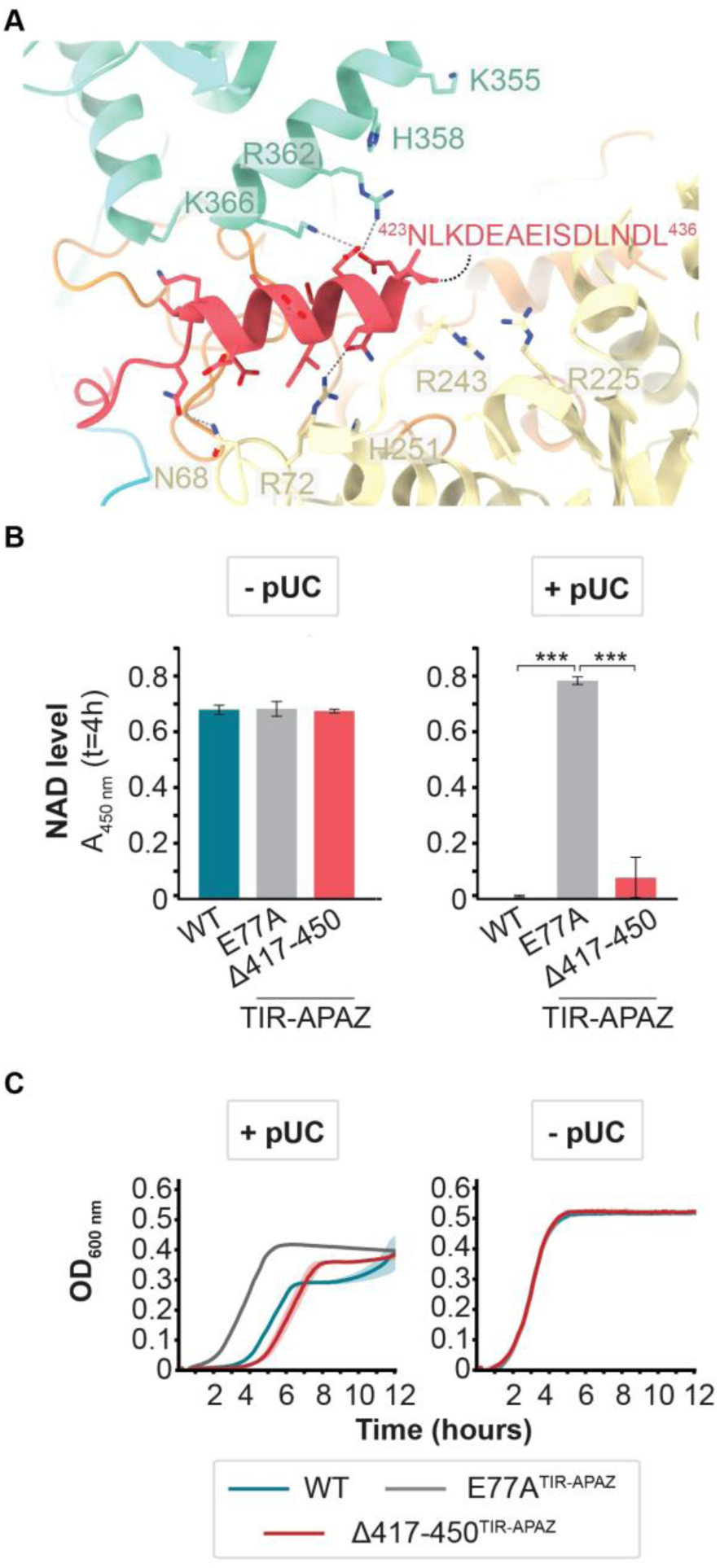
The C-terminal tail of TIR-APAZ blocks the nucleic acid binding cleft of monomeric SPARTA. (**A**) Close-up view of the interactions of the C-terminal tail with the MID and APAZ domains. (**B-C**) The C-terminal tail regulates SPARTA activity. (**B**) Total NAD (NAD^+^ + NADH) level in *E. coli* cultures expressing MapSPARTA, catalytic mutant E77A^TIR-APAZ^, or C-terminal tail truncation mutant Δ417-450^TIR-APAZ^ in the absence or presence of a highly transcribed high copy number plasmid (pUC-mRFP^ΔRBS^). (**C**) Growth curves of corresponding *E. coli* cultures expressing MapSPARTA, catalytic mutant E77A^TIR-APAZ^, or C-terminal tail truncation mutant Δ417-450^TIR-APAZ^ in the absence or presence of pUC-mRFP^ΔRBS^. The averages of three biological replicates are shown, error bars indicate standard deviations. *p < 0.05, **p < 0.01; ***p < 0.001.

In turn, the APAZ-PIWI domain interface spans 1165 Å^2^ and comprises extensive hydrophobic interactions involving numerous conserved aromatic residues (**Figure 1E**) as well as a single salt bridge involving Glu436^pAgo^ and Arg361^TIR-APAZ^. Disruption of the salt bridge and various other surface mutants had limited to no effect on SPARTA heterodimer formation, as judged by pulldown assays, indicating that individual interface residues make limited contributions to the overall interaction (**Figure 1F, G**). However, BabSPARTA complex formation was at least partially disrupted in pAgo mutants F397E^pAgo^, W421E^pAgo^, and F397E/F411E^pAgo^, as well as in TIR-APAZ mutants Y171E^TIR-APAZ^, F410E^TIR-APAZ^, and Y171E/W175E/F410E^TIR-APAZ^ (**Figure 1F, G**; **Supplementary Figure S3**). The corresponding residues in SPARTA, SPARSA, and SPARDA systems, all of which form heterodimeric complexes ^11–13^, are also strictly aromatic or hydrophobic (**Supplementary Figures S4** and **S5**). This indicates that short pAgo and TIR-APAZ proteins interact through cumulative hydrophobic/aromatic interactions and that such interactions are conserved in various short pAgos and their APAZ partners.

**Figure 3.**
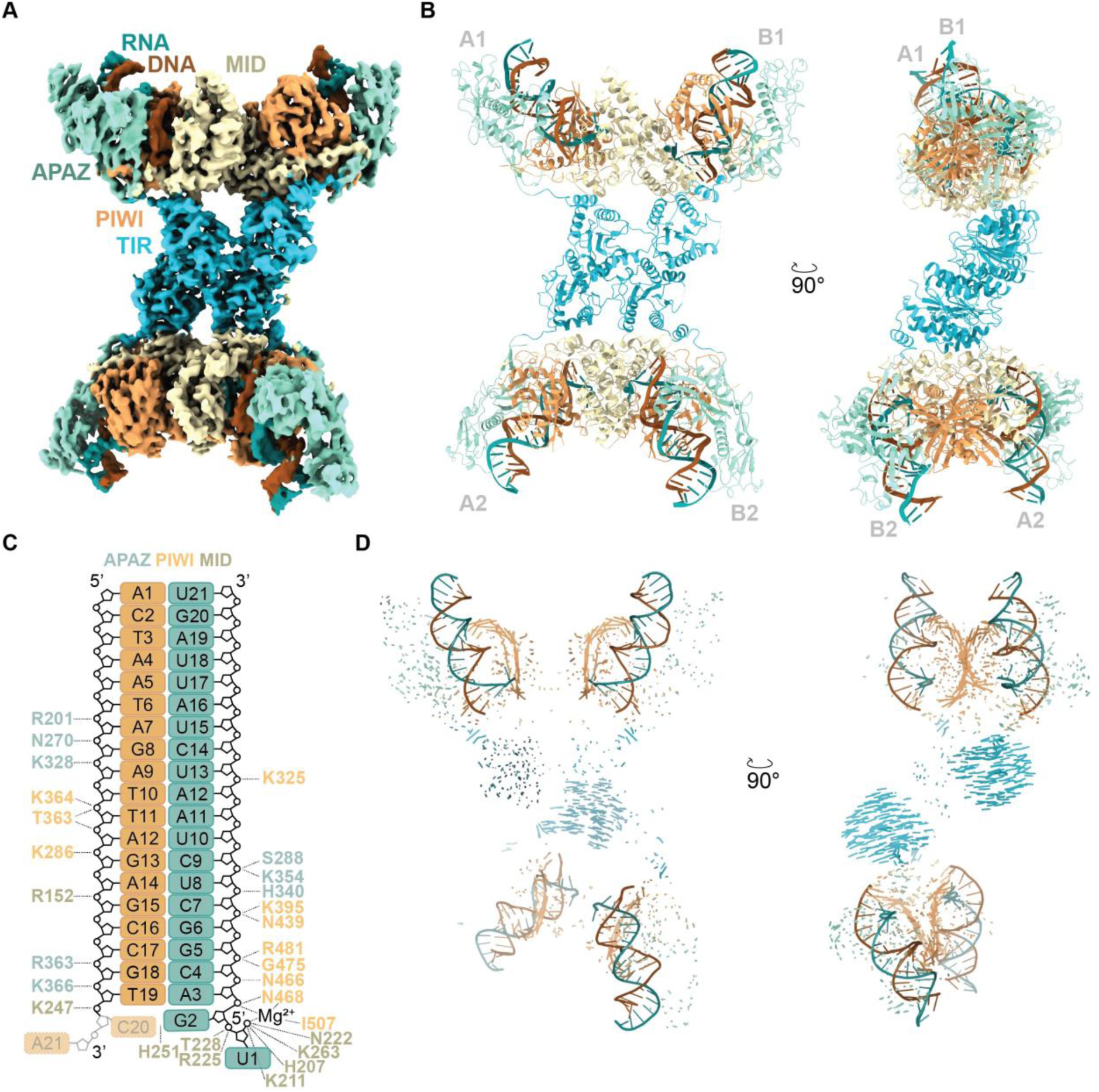
Molecular architecture of the activated tetrameric SPARTA complex. (**A**) Cryo-electron microscopy (cryo-EM) densities of the tetramer of guide RNA/target DNA duplex-bound BabSPARTA complexes. SPARTA protein domains are colored according to the scheme Figure 1A, the guide/target duplex is colored according to the scheme in panel (A). (**B**) Cartoon representation of the overall structure of the tetrameric BabSPARTA complex. (**C**) Schematic representation of the interactions of SPARTA with the guide RNA-target DNA duplex. Transparent and dotted rectangles indicate structurally disordered nucleotides. Hydrogen-bonding and electrostatic interactions are indicated with dotted lines. (**D**) Vector map displaying observed domain movement between the apo-BabSPARTA complex and each of the BabSPARTA protomers in the activated BabSPARTA complex. Vectors are colored corresponding to the domain coloring used in other panels.

**Figure 4.**
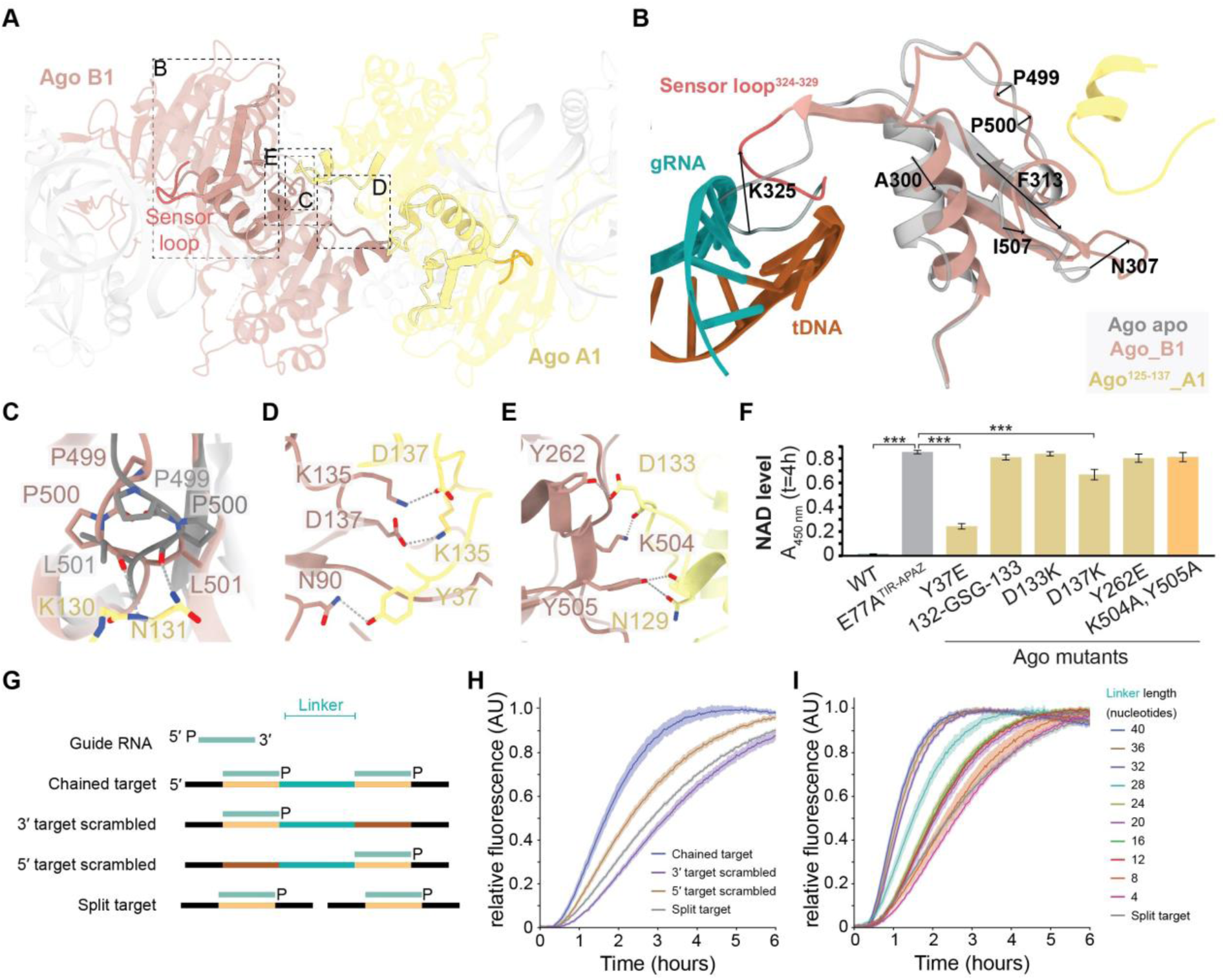
RNA-guided target DNA binding induces pAgo-pAgo interactions. (**A**) Guide RNA/target DNA duplex binding is associated with conformational changes of the sensor loop and other segments within the PIWI domain. (**B-E**) Structural rearrangements facilitate the formation of polar interactions and salt bridges between MID and PIWI domain residues. See also Supplementary Figure S10. (**F**) Mutational analyses reveal that **p**Ago-**p**Ago interaction residues are crucial for catalytic activation of SPARTA. The total NAD (NAD^+^ + NADH) level was determined in *E. coli* cultures expressing MapSPARTA, catalytic mutant E77A^TIR-APAZ^, or MapSPARTA with mutations at the pAgo-pAgo interface, in presence of a highly transcribed high copy number plasmid (pUC-mRFP^ΔRBS^). The averages of three biological replicates are shown, error bars indicate standard deviations. *p < 0.05, **p < 0.01; ***p < 0.001. See also Supplementary Figure S11. (**G-J**) SPARTA activation is increased by target sites co-localized in *cis*. BabSPARTA was mixed with guide RNA and ssDNA targets that contain a single target site (split target) or two target sites (chained target) complementary to the guide RNA, with equal abundance of binding sites in every condition (G). After addition of ε-NAD^+^ fluorescence was measured over time. SPARTA shows increased NADase activity with chained targets compared to split targets and chained targets in which one of the target sites is scrambled (H). SPARTA activation by chained targets is influenced by the linker length that connect the two target sites (I). In panel H and I, measurements are corrected for a control without ssDNA target, and normalized to the minimum and maximum of each individual reaction. The averages of three technical replicates are shown, shadings indicate standard deviations.

**Figure 5.**
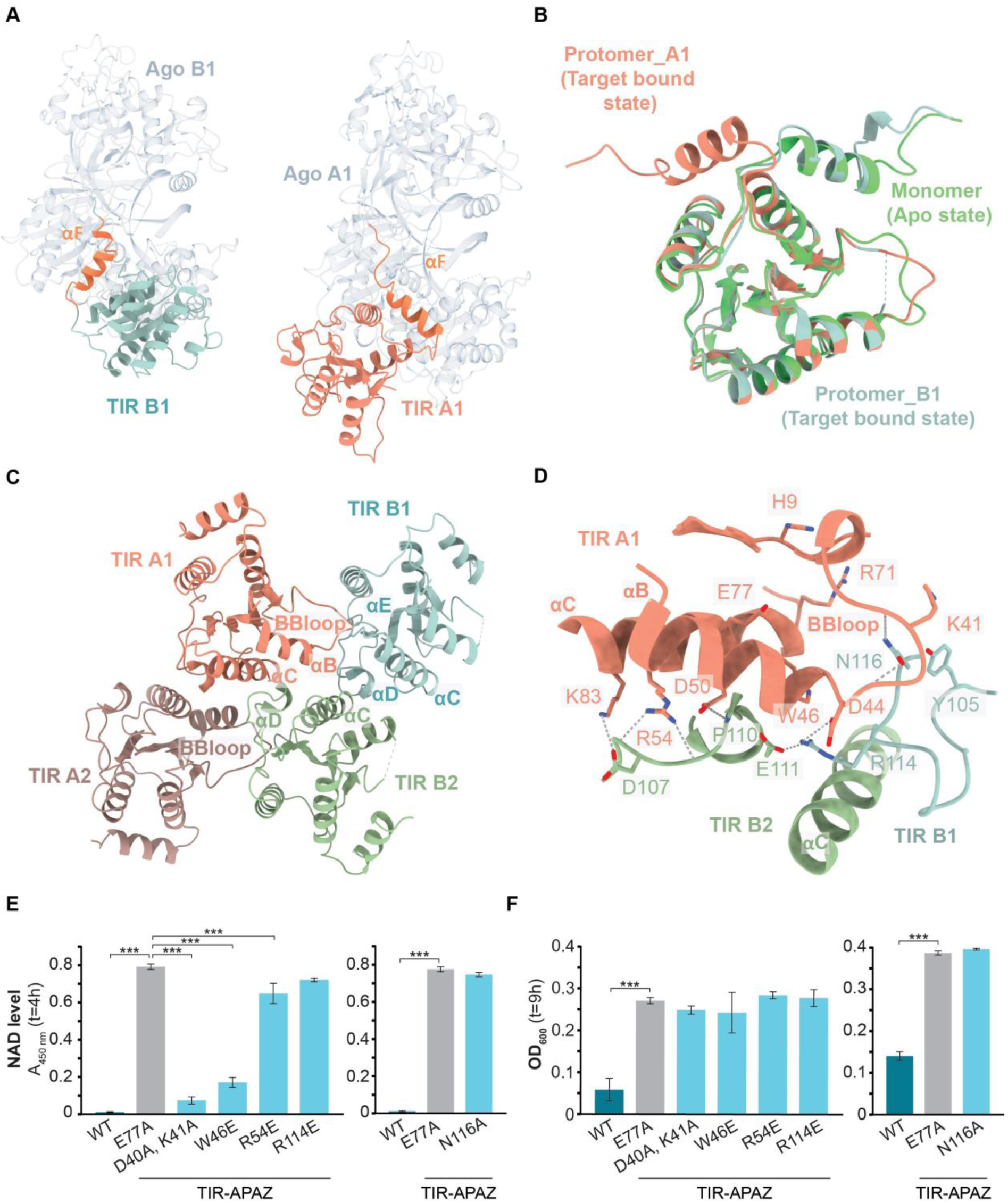
TIR domain repositioning facilitates tetramerization and catalytic activation. (**A**) Comparison of the TIR domain orientation between two SPARTA protomers interacting after tetramerization. The TIR domain in protomer A1 (in pink) is rotated 177° with respect to the TIR domain in protomer B1 (teal), with helix αF (bright orange) functioning as a hinge. Alignment performed with respect to the pAgo-APAZ moieties (grey). (**B**) Alignment of the TIR domains of the inactive apo monomeric SPARTA structure (green) with the TIR domains of two asymmetric SPARTA protomers from the activated tetrameric SPARTA structure (orange and teal). The overall TIR domain structure is maintained, except for the rotation of the αF helix (bright orange) in protomer A1 (orange) of the activated tetrameric SPARTA structure. (**C**) Overview of the assembly of the four TIR domains (TIR_A1/B1/A2/B2) in the activated tetrameric SPARTA structure. (**D**) TIR-TIR interfaces and key residues involved in TIR-TIR dimerization, TIR tetramerization, and TIR catalytic activity. (**E-F**) TIR-TIR interface residues are crucial for catalytic activation of SPARTA. The total NAD (NAD^+^ + NADH) level (E) or OD600 nm (F) was determined in *E. coli* cultures expressing MapSPARTA, catalytic mutant E77A^TIR-APAZ^, or MapSPARTA with mutations at the TIR-TIR interfaces, in presence of a highly transcribed high copy number plasmid (pUC-mRFP^ΔRBS^). The MapSPARTA and the E77A^TIR-APAZ^ mutant controls in the right panel are identical to those in Figure 1 (same experiment). The averages of three biological replicates are shown, error bars indicate standard deviations. *p < 0.05, **p < 0.01; ***p < 0.001. See also Figure S13.

### Monomeric SPARTA is autoinhibited by TIR-APAZ C-terminal tail

The relative spatial disposition of the BabTIR-APAZ and BabAgo MID and PIWI domains creates a deep, positively charged cleft that corresponds to binding site for the guide-target duplex in long pAgos (**Figure 1B**; **Supplementary Figure S1C, D** and **Supplementary Figure S7A**). Unlike long pAgos, in the monomeric BabSPARTA complex the cleft is occupied by the C-terminal tail (Ct) of TIR-APAZ (**Figure 2A**). Residues Asn423-Leu436^TIR-APAZ^ form an alpha-helix, while the remainder of the Ct (residues 436-450^TIR-APAZ^) is structurally disordered (**Figure 2A**). The Ct is highly acidic and makes numerous electrostatic contacts, via side chains of Asp426^TIR-APAZ^, Glu427^TIR-APAZ^, Glu429^TIR-APAZ^, Asp432^TIR-APAZ^, and Asp435^TIR-APAZ^, and possibly additional acidic residues in the structurally disordered remainder of the Ct (residues Asn437-As451^TIR-APAZ^), with positively charged residues in the APAZ (Lys366^TIR-APAZ^, Arg362^TIR-APAZ^) and MID (Arg72^pAgo^, and Arg225^pAgo^, Arg243^pAgo^ and His251^pAgo^) domains (**Figure 2A**). The placement of the TIR-APAZ Ct in the inactive monomeric BabSPARTA complex suggests that it may sterically inhibit guide RNA and/or target DNA binding, thus modulating SPARTA activation in an autoinhibitory manner.

To investigate the importance of the Ct in SPARTA systems, we tested the activation of SPARTA systems containing wild-type and C-terminally truncated (Ct-truncated) TIR-APAZ subunits *in vivo*. As BabSPARTA shows limited activity at 37°C (**Supplementary Figure S6**), we instead tested the activity of *Maribacter polysiphoniae* SPARTA (MapSPARTA), which displays 80% sequence identity with BabSPARTA, and Ct-truncated MapSPARTA mutants (Δ431-450^TIR-APAZ^, Δ421-450^TIR-APAZ^, and Δ417-450^TIR-APAZ^). MapSPARTA was expressed from a bacterial artificial chromosome (BAC) in the absence or presence of a highly transcribed pUC-based plasmid (pUC-mRFP^ΔRBS^). SPARTA activity was assessed by monitoring cell growth and by measuring NAD^+^ concentration in cell lysates. In the absence of invading DNA, MapSPARTA or Ct-truncated mutants thereof were not catalytically activated (**Figure 2B** and **Supplementary Figure S7B-E**). As observed previously^12^, WT MapSPARTA depleted NAD^+^ and substantially diminished the growth rate of *E. coli* in the presence of invading plasmid DNA, while MapSPARTA carrying a point mutation in the TIR domain (E77A^TIR-APAZ^) lacked NADase activity and did not affect cell growth (**Figure 2B, C** and **Supplementary Figure S7B-E**). Akin to the WT MapSPARTA, Ct-truncated mutants degraded NAD^+^ in presence of the invading pUC plasmid (**Figure 2B** and **Supplementary Figure S7C**). However, expression of the Ct-truncated mutants in the presence of invading DNA had a more severe impact on *E. coli* growth than expression of WT MapSPARTA, which suggests that the Ct-truncated mutant is hyperactive under these conditions (**Figure 2C**; **Supplementary Figure S7B-E**). Altogether, these results indicate that the Ct of TIR-APAZ is not essential for the target-dependent NADase activity of SPARTA, but that it plays an autoinhibitory role in modulating its activation mechanism *in vivo*.

**Figure 6.**
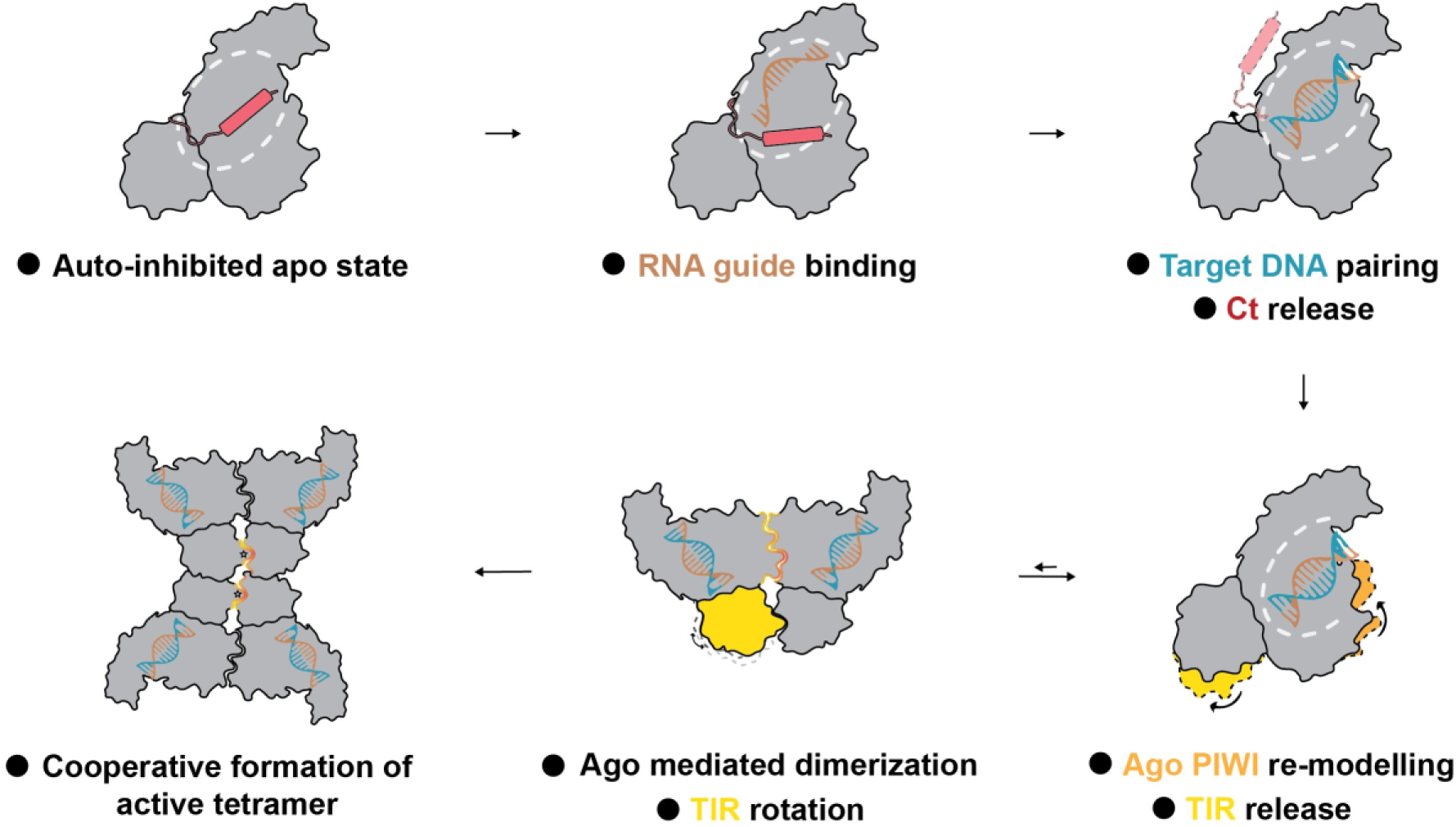
Schematic model of guide RNA-mediated target binding and catalytic activation of SPARTA.

Based on phylogeny, short pAgos can be subdivided in the S1A, S1B, S2A, and S2B clades^15^ (**Supplementary Figure S7F**). As clade S1A contains fused Sir2-APAZ-pAgo systems, we analyzed the conservation of the Ct in clades S1B (containing short pAgo/Sir2-APAZ (SPARSA) systems), S2A (containing SPARTA systems), and S2B (containing diverse short pAgo systems with various effector domains). This revealed that all APAZ domain-containing proteins in clades S2A and S2B have C-terminal extensions (on average 16-46 amino acids long) that contain a large proportion (21-45%) of negatively charged (Asp and Glu) residues, while Ct is absent in clade S1 Sir2-APAZ proteins (**Supplementary Figure S7G**). This suggests that the activity of S2-clade pAgo systems, but not that of S1 clade pAgo systems, might be regulated by C-terminal tails in the APAZ domain-containing protein.

### Molecular architecture of the activated tetrameric SPARTA complex

Guide RNA-mediated target DNA binding results in tetramerization of SPARTA complexes, which catalytically activates their TIR domains^12^. However, the mechanisms underlying tetramerization and catalytic activation of SPARTA systems are unclear. To gain structural insights into these processes, we reconstituted the active tetrameric form of BabSPARTA in the presence of a 21-nucleotide guide RNA and a complementary 21-nucleotide ssDNA, and determined its cryo-EM structure at a resolution of 3.1 Å (**Figure 3A, B**, **Supplementary Figure S8**, and **Table S1**). The reconstruction reveals a butterfly-shaped architecture formed by four BabAgo:BabTIR-APAZ SPARTA complexes, each bound to a guide RNA-target DNA duplex (**Figure 3A, B**).

The guide RNA-target DNA duplex is bound in the positively charged groove formed between the APAZ domain, and the BabAgo MID and PIWI domains, in which the negatively charged backbone phosphate groups of both the guide RNA and the target DNA are coordinated by salt-bridge and hydrogen-bonding interactions with conserved positively charged and polar residues (**Figure 3C**). The Ct of the TIR-APAZ subunit is displaced from the binding groove and structurally disordered (**Supplementary Figure S9A**), indicating conformational flexibility, in agreement with its presumed modulatory role in SPARTA activation.

As observed for other pAgo-guide RNA complexes, the 5′-terminal phosphate group of the guide RNA is sequestered in a highly conserved MID domain pocket and coordinated by a Mg^2+^ ion, which precludes base pairing with a complementary nucleotide in the DNA target strand (**Figure 3C** and **Supplementary Figure S9B**). There are no base-specific interactions with the 5′-terminal nucleotide, in agreement with the observation that guide RNAs with any 5′-terminal nucleotide mediate SPARTA activation (**Supplementary Figure S9C-E**). Unlike in other pAgos ^19, 36–38^, the guide RNA nucleotide in the second position also remains unpaired, and base pairing with target DNA is precluded by residue His251^pAgo^, which is conserved in closely related SPARTA systems (**Figure 3C** and **Supplementary Figure S4**). In line with these observations, mismatches with the target at guide nucleotide positions 1 and 2 do not impair SPARTA activation^12^.

The tetrameric assembly of the active BabSPARTA complex is a dimer of asymmetric dimers. Within each asymmetric dimer (protomers A1-B1 and A2-B2, **Figure 3A, B**), the MID, PIWI and APAZ domains are related by nearly perfect two-fold rotational symmetry, while the TIR domains adopt distinct orientations with respect to the other domains (**Figure 3D**). The orientation of one TIR domain resembles that observed in the inactive BabSPARTA monomer, while the other TIR domain is repositioned by a ∼180° rotation about a hinge segment spanning residues 143-164^TIR-APAZ^ (**Figure 3D**). The release and repositioning of the TIR domain promotes TIR-TIR interactions within each asymmetric dimer. In turn, the dimerized TIR domains of two asymmetric SPARTA-SPARTA dimers interact to form the tetrameric active SPARTA complex (**Figure 3A, B**), as detailed below.

### RNA-guided target DNA binding induces pAgo-pAgo interactions

Comparisons of the inactive monomeric and active tetrameric BabSPARTA complexes reveal conformational changes in the BabAgo subunit brought about by RNA-guided target DNA binding. In the monomeric complex, a loop spanning residues Glu324-His329^pAgo^ in the PIWI domain projects into the nucleic acid binding groove (**Figure 2C** and **Figure 4A, B**). This “sensor loop” is displaced by the guide RNA-target DNA duplex, which is accompanied by allosteric restructuring of the upstream beta hairpin (residues Ala300-Gln323^pAgo^) and an adjacent alpha helix with a downstream loop (residues Gly475-Leu501^pAgo^) in the PIWI domain (**Figure 4B** and **Supplementary Figure S10**). These structural rearrangements reshape the surface of the pAgo subunit, which facilitates pAgo-pAgo interactions and thereby SPARTA-SPARTA dimer formation. Restructuring of the Pro499-Leu501^pAgo^ segment enables BabAgo-BabAgo dimerization via the formation of hydrogen bonding contacts with the backbone carbonyl and side chain amide groups of Asn131^pAgo^ in the MID domain of the other BabAgo protomer (**Figure 4C** and **Supplementary Figure S10**). This interaction is further reinforced by additional MID-MID contacts involving salt bridges between Lys135^pAgo^ and Asp137^pAgo^, and MID-PIWI contacts involving the Asn129-Asp133^pAgo^ loop in one BabAgo protomer with residues Tyr262^pAgo^ and Lys504-Tyr505^pAgo^ in the other protomer (**Figure 4D, E**).

To verify the importance of the BabAgo-BabAgo dimerization interface residues, we investigated the effects of mutations in the corresponding residues in MapSPARTA on intracellular NAD levels and cell growth upon expression in *E. coli* in the presence of a target plasmid (pUC-mRFP^ΔRBS^) (**Figure 4F** and **Supplementary Figure S11**). MapSPARTA systems harboring mutations corresponding to Y37E^pAgo^ or D137K^pAgo^ exhibit lower NADase activities compared to WT MapSPARTA (72% and 22% reduction in NAD levels, compared to 99% reduction for WT). In turn, mutants corresponding to point mutations D133K^pAgo^ or Y262E^pAgo^, the double mutation K504A/Y505A^pAgo^, or the insertion mutation Glu132-Gly-Ser-Gly-Asp133^pAgo^ completely abolished NADase activity *in vivo* (**Figure 4F** and **Supplementary Figure S11**). For all mutants, cell growth was comparable as for cells expressing the catalytically inactive E77A^TIR-APAZ^ mutant (**Supplementary Figure S11**). These results corroborate the importance of key pAgo-pAgo interacting residues in the catalytic activation of SPARTA systems.

The BabAgo MID domain residue Lys395^pAgo^ interacts directly with both the backbone phosphate of guide RNA nucleotide 6 and BabTIR-APAZ residue Asn174^TIR-APAZ^ (**Supplementary Figure S12**). SPARTA activity is highly sensitive to mismatches at this position^12^. Despite the observation that these residues assume the same conformations in the apo and activated SPARTA complexes, we hypothesized that this interaction might be important for facilitating guide-target duplex interactions and/or contribute to SPARTA activation. Mutational analyses revealed that MapSPARTA systems harboring a mutation corresponding to K395A^pAgo^ exhibited lower NADase activities as compared to WT MapSPARTA (45% reduction in NAD levels, compared to 99% reduction for WT), while a mutation corresponding to N174A^TIR-APAZ^ completely abolished NADase activity (**Supplementary Figure S12**). For both mutants, cell growth was similar to cells expressing the catalytically inactive E77A^TIR-APAZ^ mutant. Both Lys395^pAgo^ and Asn174^TIR-APAZ^, as well as the majority of other pAgo-pAgo interface residues, are conserved in SPARTA orthologs and to some extent also in other short pAgo systems (**Supplementary Figure S4** and **S5**). This suggests that target DNA-dependent pAgo-pAgo dimerization is not only important for the activation mechanism of SPARTA, but is likely conserved in other short pAgo systems.

While the molecular architecture of the target-activated SPARTA tetramer suggests that the monomeric SPARTA complex initially dimerizes via pAgo-pAgo interactions upon binding or the target DNA, dimeric SPARTA assemblies are not readily detectable during size-exclusion chromatography^12^. This suggests that pAgo-pAgo dimers are unstable. SPARTA tetramers are observed during size-exclusion chromatography^12^, which implies that subsequent cooperative SPARTA tetramerization stabilizes the active conformation. The architecture of the SPARTA tetramer suggest that the target sites need to be located ∼90 Å apart so as to be simultaneously engaged by two SPARTA protomers within a SPARTA dimer, which would correspond to approximately 15 nucleotides. The target sites need to be located ∼150 Å apart to be simultaneously engaged by two SPARTA protomers from different SPARTA dimers, which would correspond to approximately 25 nucleotides. We hypothesized that SPARTA dimerization is enhanced upon recognition of two independent target sites located in the same DNA molecule. To test this, we designed ‘chained target’ DNAs in which two identical target sites are linked (**Figure 4G**).

We compared the NADase activity of BabSPARTA upon activation by a chained target DNA containing a 28-nucleotide linker, chained targets in which either the 5′ or the 3′ target site was scrambled, or an unchained target containing a single target site, keeping the effective concentration of target sites the same for all targets (**Figure 4G**). The chained target DNA elicited higher NADase activity as compared to the unchained target (**Figure 4H**). When either of the target sites in the chained target was scrambled, SPARTA activity remained at levels similar to that when incubated with the unchained target (**Figure 4H**). Next, we analyzed SPARTA activation using chained targets with variable linker lengths. Targets with linker lengths ≤8 nucleotides activate SPARTA at similar levels as unchained targets, suggesting that the linker is too short to stabilize SPARTA dimer formation (**Figure 4I**). In contrast, chained targets with a linker length between 12-24 nucleotides elicited higher levels of SPARTA NADase activity, and chained targets with a linker length between 32-40 nucleotides elicited even higher SPARTA NADase activity (**Figure 4I**). Corroborating our structure-based predictions, these findings suggest that targets with 12-24 nucleotide-long linkers stabilize the interactions between two SPARTA protomers within a SPARTA dimer, while targets with 32-40 nucleotide-long linkers could stabilize the interactions of SPARTA protomers within and across SPARTA dimers. Together, these results confirm that catalytic activation of SPARTA is enhanced by target sites localized in *cis*, suggesting that stabilization of SPARTA dimers and tetramers by co-localized targets is an important aspect of the activation mechanism of SPARTA.

### TIR domain repositioning facilitates tetramerization and catalytic activation

The TIR domains in the active BabSPARTA tetramer adopt two distinct orientations with respect to the corresponding BabAgo subunits (**Figure 5A, B**). In protomers B1 and B2, the orientation is nearly identical to that observed in the inactive BabSPARTA monomer. In protomers A1 and A2, however, the TIR domain and an alpha-helical segment linking the TIR and APAZ domains (residues 148-160^TIR-APAZ^) are restructured such that the TIR domain undergoes a 177° rotation with respect to the linker, centered on the hinge residue Pro147^TIR-APAZ^ (**Figure 5B**). The TIR-MID domain interactions, as formed in the monomeric SPARTA complex (**Figure 1D**) and in the tetramer protomers B1/B2, are disrupted and the TIR domain is released from the pAgo subunit. The spatial proximity of the TIR-APAZ linker to the C-terminal tail suggests that TIR domain restructuring is coupled to the displacement of the Ct upon RNA-guided target DNA binding.

The reorientation of the TIR domain in protomer A1 consequently enables the formation of interface interactions with the TIR domain in protomer B1 within the asymmetric dimer. This results in the insertion of the BB-loop ^39^ (residues As40-Trp46^TIR-APAZ^) of the rotated A1 TIR domain into a pocket in the B1 TIR domain, forming a composite NADase catalytic site capable of NAD binding (**Figure 5C, D**). The composite active site is homologous to those of other TIR domain NADases such as eukaryotic SARM1^40^ (**Supplementary Figure S13A**). Furthermore, the dimerization of the TIR domains generates an interface comprising TIR domain helices αB, αC (from the TIR domain of protomer A1) and αD (from the TIR domain of protomer B1). This interface mediates interactions with the TIR domain of protomer B2 (**Figure 5C, D**). Overall, this results in cooperative tetramerization of the TIR domains by the formation of a parallel dimer of dimers (**Figure 5C, D**).

To verify the functional significance of TIR-TIR interfaces for the activity of SPARTA systems, we tested the effects of corresponding mutations in MapSPARTA upon expression in *E. coli* in the presence of the pUC-mRFP^ΔRBS^ target plasmid (**Figure 5E, F**, and **Supplementary Figure S13A, B**). BB loop mutations D40A/K41A^TIR-APAZ^ or W46E^TIR-APAZ^ showed substantially reduced NADase activity *in vivo* (91% and 78% reductions in NAD^+^ levels, respectively, as compared to 99% reduction for WT MapSPARTA). Mutations of TIR tetramerization interface residues TIR-APAZ^R54E^ and TIR-APAZ^R114E^ abolished NADase activity completely (**Figure 5E, F**, and **Supplementary Figure S13B, C**). For all mutants, cell growth was comparable to cells expressing the catalytically inactive E77A^TIR-APAZ^ mutant, indicating loss of interference activity. Collectively, these results confirm the importance of both the TIR-TIR dimerization and the tetramerization interfaces for catalytic activation of SPARTA, and indicate that cooperative tetramerization of target-bound SPARTA complexes, mediated by TIR domain interactions, is essential for the activity of SPARTA systems.

## DISCUSSION

SPARTA systems couple short pAgo-mediated, RNA-guided target DNA recognition with TIR domain tetramerization to elicit catalytic activation that triggers NADase activity. This in turn depletes the cellular pool of NAD, leading to growth arrest and/or cell death, thereby providing population-level immunity against genetic parasites such as plasmids^12^. As such, the target sensing and activation of SPARTA systems needs to be tightly regulated to avoid spurious activation. To shed light on the target-dependent activation mechanism of SPARTA, we have performed structural analysis of BabSPARTA, determining atomic structure of the monomeric inactive and tetrameric nucleic acid-bound SPARTA complexes.

The monomeric SPARTA structure reveals that the TIR-APAZ and its associated short pAgo subunit form a heterodimeric complex through the mostly hydrophobic surfaces of their APAZ and MID-PIWI domains. As these hydrophobic interaction surfaces are generally conserved in short pAgo systems, we hypothesize that short pAgo systems from all clades adopt similar structural architectures with their respective interaction proteins. Homologous to the bilobed structure of long pAgos and eAgos^36, 37^, this forms a positively charged binding channel that can accommodate a guide RNA-target DNA duplex. Interactions between the TIR and MID domains in the apo-SPARTA complex prevent productive dimerization of the TIR domains requiring the insertion of the BB-loop from one TIR domain into the complementary pocket in the other TIR domain, thus keeping apo-SPARTA in a catalytically inactive state.

Furthermore, in the absence of a guide RNA-target DNA duplex, the negatively charged Ct of TIR-APAZ resides in the guide RNA-target DNA binding channel. As Ct and guide-target duplex binding are mutually exclusive for steric reasons, we infer that the Ct acts as a negative regulator of target binding, thereby providing an autoinhibitory mechanism within SPARTA system to avoid non-specific activation. This is in agreement with our observation that the deletion of the Ct results in hyperactivation of SPARTA *in vivo*. A negatively charged C-terminal extension is conserved in the APAZ-containing proteins associated with S2A and S2B clade short pAgo systems, but not in S1B clade short pAgo systems. Although the effector domains of the S2A (TIR) and S2B (DUF4365, Mrr-like, RecG/DHS-like) clade pAgo systems vary, this suggests that their activity is modulated through similar mechanisms. Possibly, the Ct contributes to short pAgo system fidelity and thereby prevents uncontrolled activation of these abortive infection systems.

In contrast to previously reported Ago structures, we do not observe pre-ordering of the guide RNA in the absence of target DNA binding. This suggests that guide RNAs are only transiently or flexibly bound, or that SPARTA systems recognize preformed guide-target duplexes. Alternatively, guide loading might require additional factors. For example, efficient guide DNA loading in long pAgos requires dsDNA substrates^37^, and eAGOs are typically loaded with dsRNA guide-passenger duplexes by Dicer^3^. Guide generation and loading mechanisms are currently poorly understood not only for SPARTA systems, but for pAgos in general. As loading of invader-specific guides is crucial for determining self (genome) vs. non-self (invading DNA) targeting, future research should focus on this aspect of pAgo mechanism.

The structure of the target-activated tetrameric SPARTA assembly reveals that the guide RNA becomes ordered upon binding of the target DNA strand. Akin to other pAgos, strictly conserved MID domain residues lock the phosphorylated 5′ end of the guide RNA in a pocket that precludes base pairing between guide nucleotide 1 and the target. In contrast to other Agos, also base pairing between guide nucleotide 2 and the target is precluded in SPARTA. Corroborating this observation, target DNA mismatches to guide RNA nucleotides 1 and 2 do not reduce but instead appear to increase SPARTA activity^12^. To our knowledge, precluding base pairing of the second guide nucleotide is unique for SPARTA systems and any potential functional relevance remains to be determined.

Based on our structural, biochemical and functional data, we propose a model for the activation mechanism of SPARTA (**Figure 6**). Guide RNA-dependent target DNA binding results in the displacement of the TIR-APAZ C-terminal tail. Given the proximity of the Ct and the linker segment connecting the TIR and APAZ domains, this might allosterically perturb the TIR-PIWI domain interface, thereby “unlocking” the TIR domain to enable TIR-TIR dimerization. Concurrently, binding of the guide RNA-target DNA duplex additionally triggers the displacement of a ‘sensor loop’ located in the PIWI domain. Restructuring of the sensor loop in turn allosterically induces further changes in the PIWI domain that result in the formation of an interface that facilitates pAgo-pAgo dimerization. In the SPARTA-SPARTA dimer, the TIR-domain of one SPARTA protomer is rotated and forms a BE helix interface with the TIR domain of the other SPARTA protomer which remains in the ‘inactive’ conformation. This TIR-TIR dimerization results in the formation of a composite catalytic site by insertion of one BB-loop into the other TIR domain. However, through formation of a BCD interface, the TIR dimer also interacts with the TIR-TIR dimer of another SPARTA dimer, forming a tetrameric assembly in a parallel configuration (**Figure 6**). Consequently, SPARTA activation requires a total of four activated SPARTA complexes forming a complex with a 4:4:4:4 stoichiometry (TIR-APAZ:pAgo:RNA:DNA). As formation of stable dimeric SPARTA complexes is not observed upon target DNA binding^12^, we conclude that TIR tetramerization is a fast, cooperative process that is essential to stabilize the initially formed SPARTA-SPARTA dimers, while the initial dimerization may be the rate limiting step. In this way, tetramerization contributes to the sensitivity, fidelity, and spatiotemporal control of SPARTA activity.

TIR domain oligomerization is a general activation mechanism concept in immune systems that use TIR as an effector domain for NADase activity or signaling ^23, 25, 28, 31, 33, 34^. However, TIR domains have distinct oligomerization configurations in diverse immune systems (**Figure 7**). For example, enzymatic TIR domains from animal SARM1, plant NLR-RPP1, and bacterial TIR-STING immune proteins arrange a double antiparallel TIR filament configuration^33, 34, 40^ in which a catalytically active filament interacts though the BE-interfaces, while the other filament forms distinct interfaces and mostly acts to stabilize the catalytic filament through AE-interfaces. On the other hand, bacterial TIR-SAVED proteins form a single enzymatic TIR filament stabilized by cyclic oligoadenylate-dependent polymerization of the SAVED domains^28^. In contrast to these TIR domains, SPARTA TIR dimers assemble in a parallel, head-to-tail orientation through a BCD interface analogous to the enzymatic TIR domain from the bacterial virulence factor Tir and scaffolding TIR oligomers formed by animal MAL and MyD88 proteins^41–43^. In SPARTA, further scaffolding-like polymerization of TIR is not possible due to steric hindrance by the pAgo subunit, which keeps the second TIR in an inhibited conformation. Overall, these structural comparisons highlight the versatility of TIR oligomerization which, despite highly variable underlying interactions, is essential for TIR functions in innate immunity in all domains of life.

**Figure 7.**
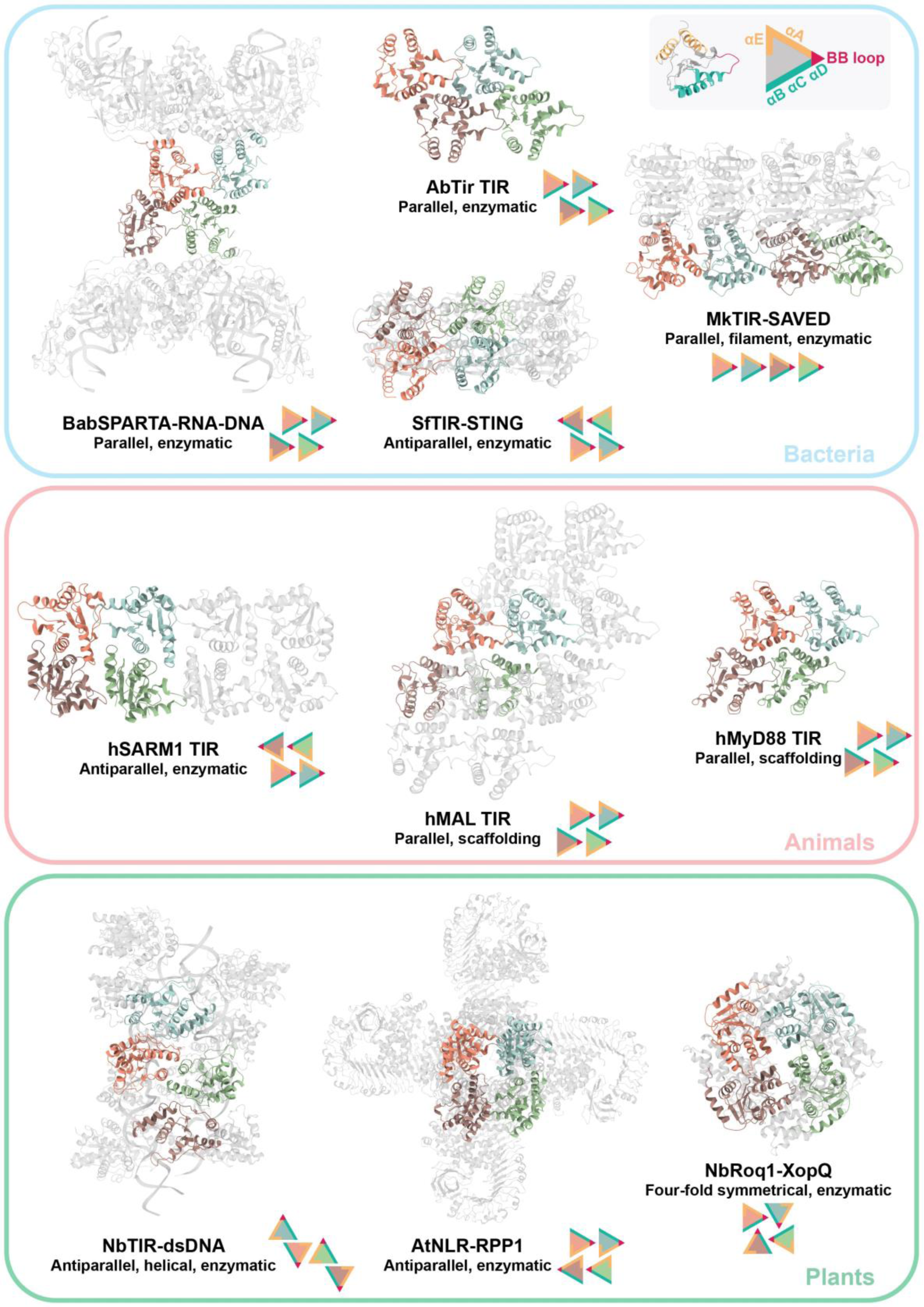
Comparison of oligomerization modes of enzymatic and scaffolding TIR domains found in bacteria, animals, and plants. Representative structures of TIR containing proteins with different origin, assembly geometry and function. Schematics of the main interaction interfaces are represented using triangles of different colors for different TIR-helices positions. (AbTir TIR, PDB: 7UXU; SfTIR-STING, PDB: 7UN8; MkTIR-SAVED, PDB: 7QQK; hSARM1 TIR, PDB: 7NAK; hMAL TIR, PDB: 5UZB; hMyD88 TIR, PDB: 6I3N; NbTIR-dsDNA, PDB: 7X5K; AtNLR-RPP1, PDB: 7CRC; NbRoq1-XopQ, PDB: 7JLV).

While this manuscript was in preparation, additional studies reporting structures of inactive and target-activated SPARTA complexes from *Crenotalea thermophila* (82% sequence ID with BabSPARTA) and *Maribacter polysiphoniae* (80% sequence ID with BabSPARTA) were published that generally corroborate our findings ^44–50^. Beyond insights derived in our study, these studies reveal structures of SPARTA bound to the guide RNA-target DNA duplex prior to oligomerization^45^ and in the dimeric form prior to tetramerization^44, 46, 50^. Furthermore, mutational analyses in these studies extend insights into the structural determinants of guide RNA and target DNA binding^46^, TIR-TIR interactions^44–47^, interactions of the TIR domains with the NAD substrate^47, 48^, as well as the importance of the Ct^44, 46^. Together with our structural analysis, these studies provide a comprehensive understanding of SPARTA mechanisms.

Finally, as structure-guided mutants have improved the activity and fidelity of CRISPR-Cas-based nucleic acid detection and genome editing tools^51–54^, we expect that this work provides a framework for further development of SPARTA-based nucleic acid detection tools^12^. In conclusion, this study provides critical insights into the structural architecture of SPARTA systems and the molecular mechanisms that control the catalytic activation of their TIR domains. It contributes to our understanding of the general mechanisms underlying the biological function of short pAgo systems, and enables their further exploitation as programmable molecular tools.

## DATA AVAILABILITY

Plasmid and oligonucleotide sequences, multiple sequence alignments, C-terminal tail sequence analysis, and raw data underlying bar graphs will be made available via Mendeley Data.

The protein expression plasmid encoding MapSPARTA (pBK086) and BabSPARTA (pAP007) have been deposited on Addgene (Plasmid #183145 and #206809). Other plasmids are available upon request.

Atomic coordinates and cryo-EM maps have been deposited in the protein data bank (PDB entry ID 8QLO and 8QLP) and Electron Microscopy Data Bank (EMDB, entry ID EMD-18486 and EMD-18487).

## ACKNOWLEDGEMENTS

We thank members of the Swarts and Jinek labs for discussion.

## AUTHOR CONTRIBUTIONS

G.F., B.K., M.J., and D.C.S. designed experiments. G.F. and M.J. determined and analyzed SPARTA structures. B.K., and A.P., purified proteins and B.K., A.P., C.H., and D.C.S. performed and/or analyzed biochemical and *in vivo* experiments. G.F., B.K., M.J., and D.C.S. wrote the manuscript and prepared figures. All authors have read and approved the manuscript.

## MATERIALS AND METHODS

### Plasmid availability

MapSPARTA expression vector (pBK086) and BabSPARTA expression vector (pAP007) have been deposited to AddGene (Plasmid #183145 and #206809). Other plasmids are available upon request.

### Plasmid cloning for *in vivo* assays and protein production

All cloning procedures were performed as described previously^1^ The genes encoding BabSPARTA (Coding contig: JADBKC010000166.1; BabAgo: MBE3571069.1.1; BabTIR-APAZ: MBE3571068.1) were ordered from Twist Bioscience as an artificial operon. This operon was cloned in the pET-His6-MBP-TEV-LIC cloning vector. Bacterial Artificial Chromosomes (BACs) encoding MapSPARTA or the catalytically inactive mutant E77A^TIR-APAZ^ were previously constructed and described^1^ All plasmids used in this study will be listed on Mendeley Data.

### SPARTA protein expression and purification

Proteins were purified as described previously^1^.

### Cryo-EM sample preparation and data collection: BabSPARTA-gRNA

Purified BabSPARTA was mixed with a 5′-phosphorylated RNA guide (oBK084, 5′-UGACGGCUCUAAUCUAUUAGU-3’) in assembly buffer (20 mM HEPES pH 7.5, 125 mM KCl, 2 mM MgCl_2_). The final sample contained 20 μM BabSPARTA and 30 μM gRNA (1:1.5 molar ratio) in a total volume of 80 μL. The volume was incubated at 50 °C for 1 hour, centrifuged at 18000 rpm for 10’ at room temperature and directly used for cryo-EM grid preparation.

3.5 µL of sample were applied to a freshly glow discharged 200-mesh Au R1.2/1.3 grid (Quantifoil Micro Tools), incubated for 5 s, blotted for 6 s at 100% humidity, 4 °C, plunge frozen in liquid ethane (using a Vitrobot Mark IV plunger, FEI) and stored in liquid nitrogen. Cryo-EM data collection was performed on a FEI Titan Krios G3i microscope (University of Zurich, Switzerland) operated at 300 kV and equipped with a Gatan K3 direct electron detector in super-resolution counting mode. A total of 14332 movies were recorded at 130000x magnification, resulting in a super-resolution pixel size of 0.325 Å. Each movie comprised 36 subframes with a total dose of 58.005 e^-^/Å^2^. Data acquisition was performed with EPU Automated Data Acquisition Software for Single Particle Analysis (ThermoFisher Scientific) with three shots per hole at -1.0 mm to -2.4 mm defocus (0.2 mm steps).

### Data processing and model building: BabSPARTA-gRNA

The collected exposures were processed in cryoSPARC (v.4.2)^55^. Patch Motion Correction and Patch CTF Correction were used to align and correct the imported 14332 movies. Blob picker (minimum particle diameter 90 Å, maximum particle diameter 180 Å) was used to select particles that were further extracted (extraction box size 360 pix, Fourier-cropped to box size 120 pix) and classified in 50 classes using 2D Classification. 22 classes, corresponding to 2815212 particles, were selected and given as input to a 2-classes Ab-Initio Reconstruction. One of the two classes, corresponding to 1733366 particles, was further classified using a 5-classes Heterogeneous Refinement. One of the 5 classes, corresponding to 904721 particles, was used as input for Non-uniform Refinement, resulting in a 2.73 Å (GSFSC resolution) density. A summary of the processing workflow is shown in the Supplementary Figure S6.

An initial model of BabSPARTA was generated using AlphaFold2 ColabFold^56^. The model was manually docked as rigid body in the BabSPARTA-gRNA cryoEM density map using UCSF ChimeraX^57^, followed by real space fitting with the Fit in Map function. The model was subjected to manual refinement against the corresponding cryoEM map using the software Coot^58^ and real space refine in Phenix^59^. Secondary structure restraints, side chain rotamer restraints and Ramachandran restraints were used. The refinement statistics will be listed on Mendeley Data. The final model comprises one copy of BabTIR-APAZ^1–435^ and one copy of BabAgo^1–151, 204–507^. Partial density for nucleotide 2 of the RNA guide was visible in the map, but not interpretable, probably due to partial occupancy. The molecule was therefore not modelled. Figures preparation of model and map was performed using UCSF ChimeraX^57^.

### Cryo-EM sample preparation and data collection: BabSPARTA-gRNA-tDNA

Purified BabSPARTA was mixed with a 5′-phosphorylated guide RNA (oBK084, UGACGGCUCUAAUCUAUUAGU) in assembly buffer (20 mM HEPES pH 7.5, 125 mM KCl, 2 mM MgCl_2_). The mixture was incubated at 37 °C for 30’ before adding the target DNA (oBK382, CAACTAATAGATTAGAGCCGTCAAT). The final sample contained 14.5 μM BabSPARTA, 22.5 μM gRNA and 22.5 μM tDNA (1:1.5:1.5 molar ratio) in a total volume of 100 μL. The volume was incubated at 50 °C for 1 hour, centrifuged at 18000 rpm for 10’ at room temperature and analyzed by size exclusion chromatography (Superdex 200 Increase 15/500 GL column, ÄKTA pure™ micro system) (Supplementary figure S5, Panel B). The fractions corresponding to the peak at the elution volume ∼1.3 mL were pooled together (total volume 300 µL) and concentrated to 30 µL using a 100000 Da molecular weight cut-off centrifugal filter (Merck Millipore). n-Octyl-β-D-Glucoside was freshly added to the sample to a final concentration of 0.07% m/V.

2.4 µL of sample were applied to a freshly glow discharged 200-mesh Cu + 2 nm C R1.2/1.3 grid (Quantifoil Micro Tools), blotted for 6 s at 100% humidity, 4 °C, plunge frozen in liquid ethane (using a Vitrobot Mark IV plunger, FEI) and stored in liquid nitrogen. Cryo-EM data collection was performed on a FEI Titan Krios G3i microscope (University of Zurich, Switzerland) operated at 300 kV and equipped with a Gatan K3 direct electron detector in super-resolution counting mode. A total of 9041 movies were recorded at 130000x magnification, resulting in a super-resolution pixel size of 0.325 Å. Each movie comprised 47 subframes with a total dose of 59.347 e^-^/Å^2^. Data acquisition was performed with EPU Automated Data Acquisition Software for Single Particle Analysis (ThermoFisher Scientific) with three shots per hole at -1.0 mm to -2.4 mm defocus (0.2 mm steps).

### Data processing and model building: BabSPARTA-gRNA-tDNA

The collected exposures were processed in cryoSPARC (v.4.2)^55^. Patch Motion Correction and Patch CTF Correction were used to align and correct the imported 9041 movies. Blob picker (minimum particle diameter 100 Å, maximum particle diameter 300 Å) was used to select particles that were further extracted (extraction box size 640 pixel, Fourier-cropped to box size 128 pixel) and classified in 50 classes using 2D Classification. 18 classes, corresponding to 865048 particles, were selected and given as input to a 2-classes Ab-Initio Reconstruction. One of the two classes, corresponding to 486647 particles, was refined with Non-uniform Refinement. As some parts of the map showed signs of flexibility, particles and mask were used as input for a 3D variability analysis, solving 3 modes and using 5.5 Å as filter resolution. The results were analyzed with 3D Variability Display in cluster output mode with 5 clusters. One of the 5 cluster volumes, corresponding to 60319 particles, was used as input for Non-uniform Refinement, resulting in a 3.09 Å (GSFSC resolutionn) density. A summary of the processing workflow is shown in the Supplementary Figure S7.

Four copies of the BabSPARTA apo model were manually docked as rigid bodies in the cryoEM density map of BabSPARTA-gRNA-tDNA using UCSF ChimeraX^57^, followed by real space fitting with the Fit in Map function. The DNA and RNA molecules were manually built using the software Coot^58^. The model was subjected to manual refinement in Coot^58^ against the corresponding cryoEM map and real space refine in Phenix^59^. Secondary structure, side chain rotamer, Ramachandran and nucleic acid restraints (base pair, stacking plane and sugar pucker restraints calculated with the LibG^60^ script) were used. The refinement statistics will be listed on Mendeley Data. The final model comprises 4 copies of BabTIR-APAZ^1–416^, 4 copies of BabAgo^1–152, 204–289, 294–507^, four copies of the duplex RNA^1–20^-tDNA^3–20^ and 4 Mg^2+^ ions. Figures preparation of model and map was performed using UCSF ChimeraX^57^.

### Sequence conservation analysis

Short pAgo and APAZ-domain containing proteins sequences were obtained from previous analyses and alignments described previously^2^ and extended with sequences from (BabAgo (MBE3571069.1) and BabTIR-APAZ (MBE3571068.1) and NgaSPARDA (NgaAgo and NgaDUF4365-APAZ)^3^. In total 285 short pAgos an 255 APAZ-domain containing proteins were used for multiple sequence alignments. Sequences were aligned in Jalview (V2.11.2.7) using MAFFT^4^ with E-INS-I settings. Full sequence alignments can will be listed on Mendeley Data. For data visualization, the aligned sequences were limited to a subset of proteins, gaps were hidden, and sequence conservation was visualized together with structural features from the heterodimeric BabSPARTA complex using ESpript 3.0 (https://espript.ibcp.fr).

For C-terminal tail conservation analysis the sequence alignment of clade S1A, S2A, and S2B APAZ-domain was used. All (aligned) sequences up to BabTIR-APAZ residue 416 were removed (i.e. only sequences aligned with the sequence downstream of BabTIR-APAZ residue 417 were included). Of these sequences, length and Asp/Glu content was determined in Excel. C-terminal tail sequences and analysis will be listed on Mendeley Data.

### ε-NAD assays

To assess NADase activity *in vitro* we followed protocols described previously^1^. A reaction mixture of purified BabSPARTA complex in SEC buffer, ε-NAD^+^ (Nicotinamide 1,N6-ethenoadenine dinucleotide), RNA guide, and 5X reaction buffer (50 mM MES pH 6.5, 375 mM KCl, and 10 mM MgCl_2_) was prepared on ice in 96-well or 384-well plates. The mixture was incubated at room temperature for 15 min, after which DNA target was added. Unless indicated otherwise, concentrations of each component were 1 µM SPARTA complex, 1 µM guide, 25 µM ε-NAD^+^, 10 mM MES pH 6.5, 125 mM KCl, and 2 mM MgCl_2_ in a final volume of 60 µL or 20 µL. In the assays to determine the preferred 5′-end nucleotide of the guide, the target concentration was 200 nM, whereas in the assay to determine the temperature optimum of BabSPARTA the target concentration was 1 µM. For chained target experiments, chained target concentration was 100 nM, while the concentration of 5′ or 3′ scrambled chained target as well as the single target was 200 nM. After addition of the target, the plate was transferred to a SH1M2FG plate reader (Biotek) preheated to 55 °C. For kinetic measurements, fluorescence intensity was measured in kinetic mode using an excitation wavelength of 310 nm and emission wavelength of 410 nm. To determine the temperature optimum of BabSPARTA, reaction mixtures were prepared in PCR strips, and after addition of the target the reactions were transferred to a pre-heated PCR block. After 1h incubation, the reactions were transferred to a 384-well plate and the fluorescence intensity was measured as described for kinetic measurements.

The sequences of all guide and target oligonucleotides used will be listed on Mendeley Data. All measurements were corrected with a control without ssDNA target. For assays with chained targets, fluorescence values were normalized to the minimum and maximum of each individual reaction. All assays were performed as technical triplicates, error bars in bar charts and shaded areas in line graphs indicate standard deviations.

### *In vivo* total NAD measurement and growth assays

To assess the effect of MapSPARTA expression (from bacterial artificial chromosome (BAC) pBK138) and mutants thereof (pBK261-pBK273, pBK295-297, and pBK301) on total NAD concentration in *E. coli*, BW25113(DE3) we followed the protocol described by Koopal *et al*. (2022)^1^. To activate MapSPARTA, a pUC-based high copy plasmid that encodes mRFP under control of a T7 promoter (pBK145) was included. 4h after induction of MapSPARTA expression the NAD concentration was determined, and cultures were diluted 1 : 200 in 96-wells plates in LB containing 100 µg/mL arbenicillin, 25 µg/mL chloramphenicol, 0.2% arabinose, and 0.25 mM IPTG. The plate was covered with a gas permeable moisture seal (4titude) and was transferred to a Synergy Neo2 or SH1M2FG plate reader (Biotek) preheated to 37 °C. Absorbance at 600 nm was measured in kinetic mode for 12 hours. All experiments were performed as biological triplicates, error bars in bar charts and shaded areas in line graphs indicate standard deviations.

### Protein pulldown of BabAgo using BabTIR-APAZ

6x-his-MBP-tagged BabSPARTA (pAP007) or mutants thereof (pBK274-pBK290) were transformed in *E. coli* BL21 Star (DE3). Single colonies were used to inoculate 1 mL LB cultures which were incubated o/n at 37 °C at 180 RPM. The o/n cultures were used to inoculate 10 mL LB medium which were incubated at 37 °C at 180 RPM. When an OD_600_ nm of ∼0.6 was reached, the cultures were incubated on ice for 10 minutes while the temperature of the incubator was lowered to 18 °C. Protein expression was induced by addition of 0.25 mM IPTG, and cultures were further incubated at 18 °C and 180 RPM for 16 hours. Cells were harvested by centrifugation at 3500 x g at 4 °C for 15 min. Cell pellets were resuspended in 0.5 mL amylose wash buffer (500 mM NaCl, protease inhibitors (200 µg/mL AEBSF, 1 µg/mL pepstatin), 20 mM HEPES pH 7.5), transferred to thin-walled 0.5 mL Eppendorf tubes, and lysed in a sonicator bath at 4 °C (QSONICA Q700A-220 sonicator; amp 90%, 1s ON/4s OFF, 20 minutes total ON time). The lysates were centrifuged for 10 min at 20,000 x g at 4 °C. Clarified lysates were incubated for 1 hour at 4 °C on a nutator with 40 µL amylose resin slurry (New England Biolabs) equilibrated with amylose wash buffer. After incubation, the resin of each sample was pelleted by centrifugation for 30 seconds at 2000 x g at 4 °C, and resuspended in 1 mL fresh amylose wash buffer. The wash step was repeated 3 times. Finally, the resin of each sample was resuspended in 40 µL SDS-PAGE loading buffer and incubated for 5 minutes at 95 °C. The samples were analyzed on SDS-PAGE using CBB staining, and the bands corresponding to MBP-BabTIR-APAZ and BabAgo were quantified using GelAnalyzer (version 19.1; www.gelanalyzer.com).

### Statistical analyses

For *in vivo* total NAD measurements, as well OD_600 nm_ measurements after 5 hours of growth during the growth assays, statistical analyses were performed in R version 4.1.0. The data in each experiment was subjected to a Shapiro-Wilk test on linear model residuals and Levene’s test to confirm normal distribution of the data and homogeneity of variances, respectively. One-way ANOVA was used to test for significant effect of MapSPARTA variant on total NAD and OD_600 nm_, and Tukey’s test was used for post-hoc pairwise comparisons.

In three experiments the assumption of normality was not met, which was solved for two experiments by using a x^1^.^25^ transformation on the data. This concerns the statistical tests described in Figure 1, Figure 4, Figure S7 and Figure S11. For one experiment transforming the data was not successful (comparison of OD_600 nm_ between cells expressing MapSPARTA, E77A^TIR-APAZ^ and Δ417-450^TIR-APAZ^, no pUC-mRFP^ΔRBS^; see Figure S7). Therefore, instead of a one-way ANOVA a Kruskal-Wallis rank sum test was used. As this test was not significant, no post-hoc test was performed for this experiment.

## SUPPLEMENTARY DATA

**Table S1.** Cryo-EM data collection, refinement and validation statistics.

|  | BabSPARTA<br>(EMDB-18486)<br>(PDB 8QLO) | BabSPARTA-RNA-DNA<br>(EMDB-18487)<br>(PDB 8QLP) |
| --- | --- | --- |
| **Data collection and processing** |  |  |
| Magnification | 130,000 | 130,000 |
| Voltage (kV) | 300 | 300 |
| Electron exposure (e<sup>-</sup>/Å<sup>2</sup>) | 58.005 | 59.347 |
| Defocus range (μm) | -1.0 to -2.4 (0.2 steps) | -1.0 to -2.4 (0.2 steps) |
| Pixel size (Å) | 0.325 | 0.325 |
| Symmetry imposed | C1 | C1 |
| Initial particle images (no.) | 4,943,576 | 2,448,462 |
| Final particle images (no.) | 907,847 | 60,319 |
| Map resolution (Å) | 2.57 | 3.09 |
| FSC threshold | 0.143 | 0.143 |
| Map resolution range (Å) | 2.3-3.9 | 2.5-7.0 |
| **Refinement** |  |  |
| Initial model used (PDB code) | AlphaFold2 | AlphaFold2 |
| Model resolution (Å) | 2.6 | 3.1 |
| FSC threshold | 0.143 | 0.143 |
| Model resolution range (Å) | 2.5-2.9 | 3.0-3.7 |
| Map sharpening *B* factor (Å<sup>2</sup>) | -102.9 | -43.2 |
| Model composition |  |  |
| Non-hydrogen atoms | 7282 | 31660 |
| Protein residues | 892 | 3463 |
| Nucleotide residues | 0 | 168 |
| Ligands | 0 | MG: 4 |
| *B* factors (Å<sup>2</sup>) min/max/mean |  |  |
| Protein | 72.50/216.62/133.01 | 60.38/232.57/117.08 |
| Nucleotide | --- | 82.06/269.71/157.35 |
| Ligand | --- | 86.98/97.76/92.37 |
| R.m.s. deviations |  |  |
| Bond lengths (Å) | 0.009(5) | 0.005(2) |
| Bond angles (°) | 1.181(38) | 0.752 (96) |
| Validation |  |  |
| MolProbity score | 2.23 | 2.02 |
| Clashscore | 12.71 | 13.82 |
| Poor rotamers (%) | 2.03 | 0.53 |
| Ramachandran plot |  |  |
| Favored (%) | 95.37 | 94.60 |
| Allowed (%) | 4.51 | 5.40 |
| Disallowed (%) | 0.11 | 0.00 |

**Supplementary Figure S1.**
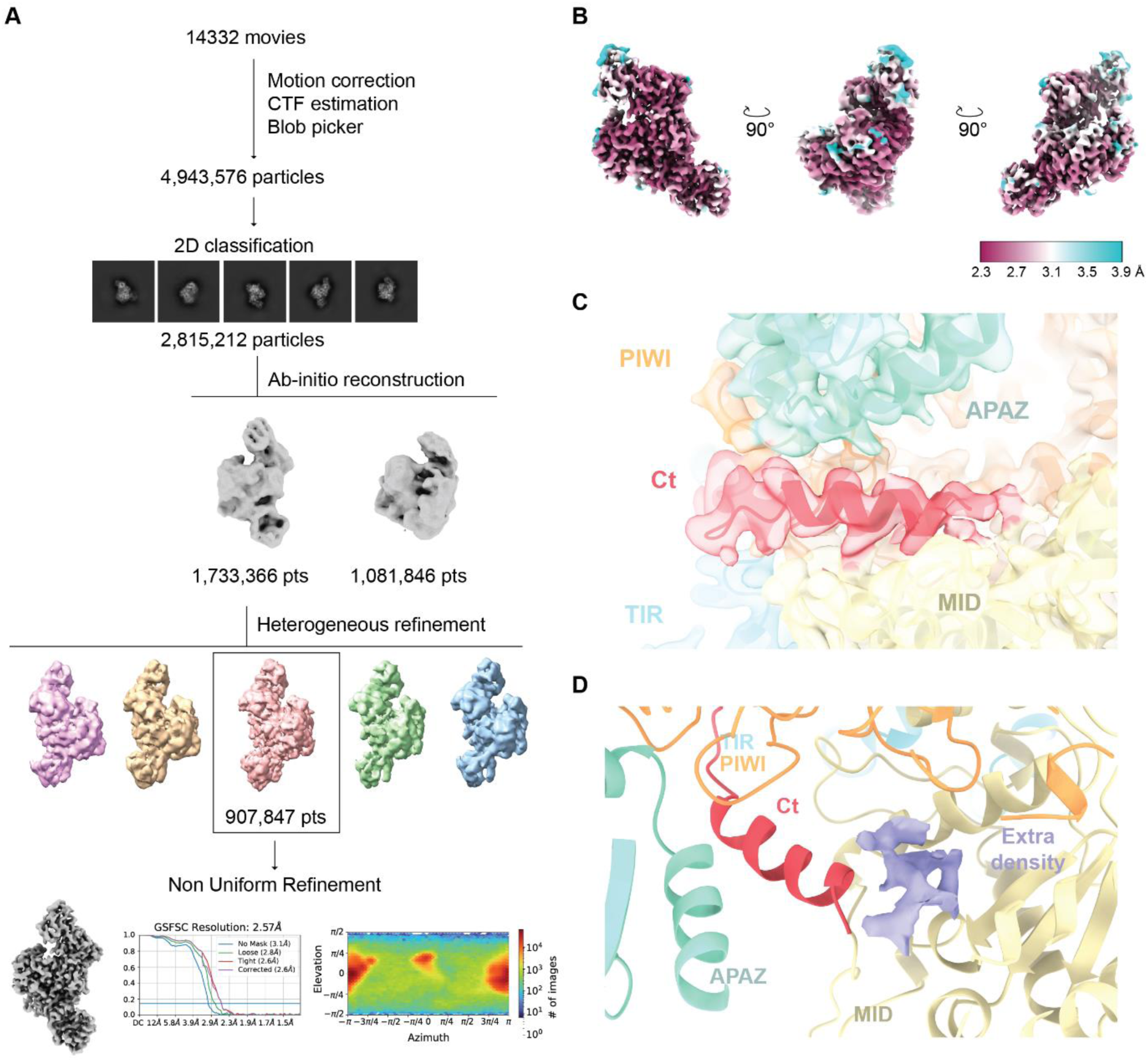
Cryo-EM data processing for BabAgo/BabTIR-APAZ complex. (**A**) Cryo-EM image processing workflow for BabAgo/BabTIR-APAZ complex. (**B**) Cryo-EM densities of the BabSPARTA complex colored according to local resolution. (**C**) Zoomed-in view of the C-terminal tail region of BabAgo/BabTIR-APAZ complex. Model displayed in cartoon representation; density depicted as transparent surface representation. Domains are colored according to the scheme in Figure 1A. (**D**) Zoomed-in view of the MID/C-terminal tail interface of BabAgo/BabTIR-APAZ complex. Model displayed in cartoon representation; an extra-density that could not be fitted with the protein model is displayed in solid surface representation (in lilac).

**Supplementary Figure S2.**
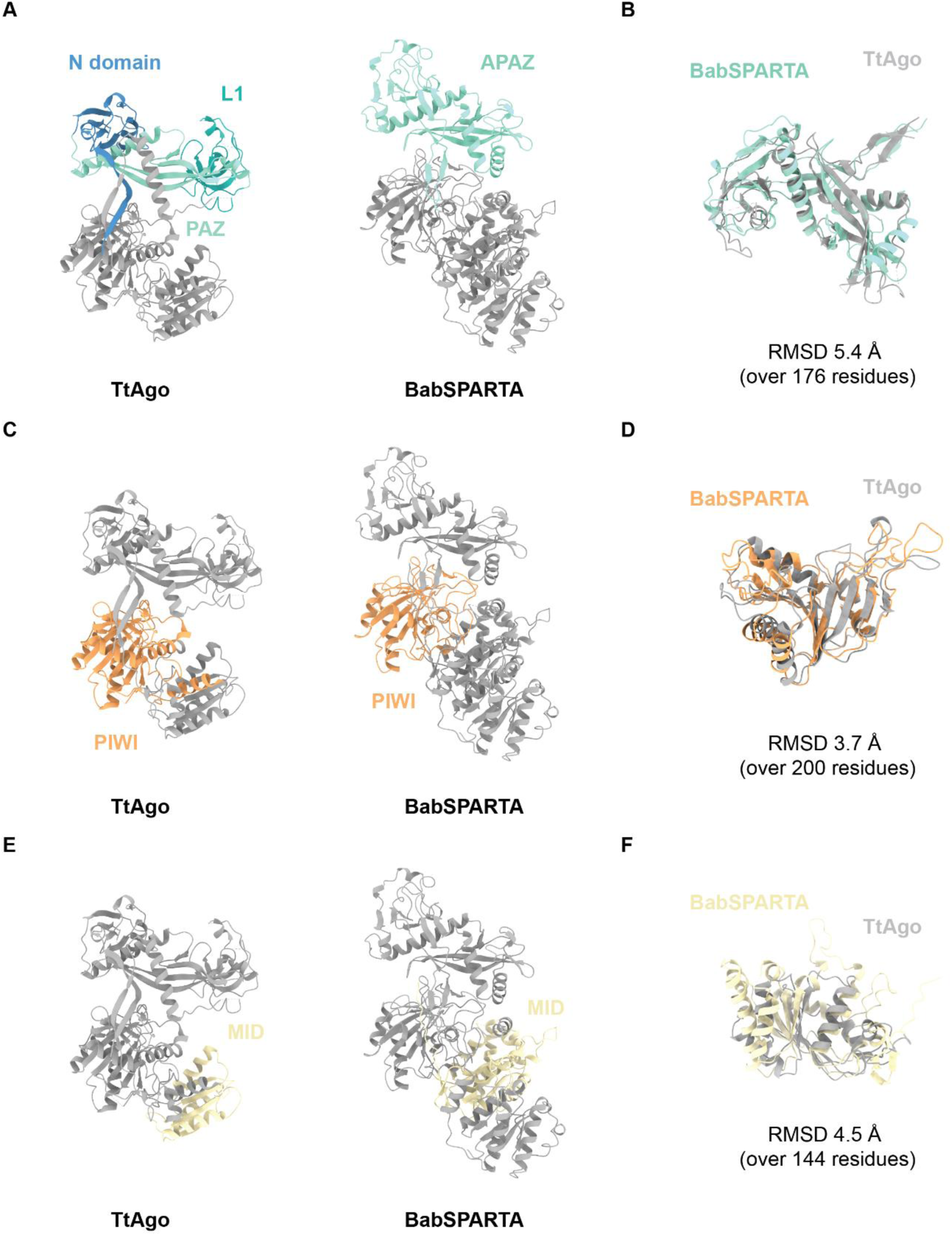
BabSPARTA shows a similar fold to canonical long-A pAgo proteins. (**A**) TtAgo and BabSPARTA structures in ribbon representation with N1-L1-PAZ domains and APAZ domain highlighted in shades of blue. (**B**) Alignment of the TtAgo N1-L1-PAZ domains (in grey, ribbon representation) with BabSPARTA APAZ domain (in blue, ribbon representation) performed with PyMOL^1^, with respective RMSD. (**C**) TtAgo and BabSPARTA structures in ribbon representation with PIWI domains highlighted in orange. (**D**) Alignment of the TtAgo APAZ domain (in grey, ribbon representation) with BabSPARTA APAZ domain (in orange, ribbon representation) performed with PyMOL^1^, with respective RMSD. (**E**) TtAgo and BabSPARTA structures in ribbon representation with MID domains highlighted in yellow. (**F**) Alignment of the TtAgo MID domain (in grey, ribbon representation) with BabSPARTA MID domain (in yellow, ribbon representation) performed with PyMOL^1^, with respective RMSD.

**Supplementary Figure S3.**
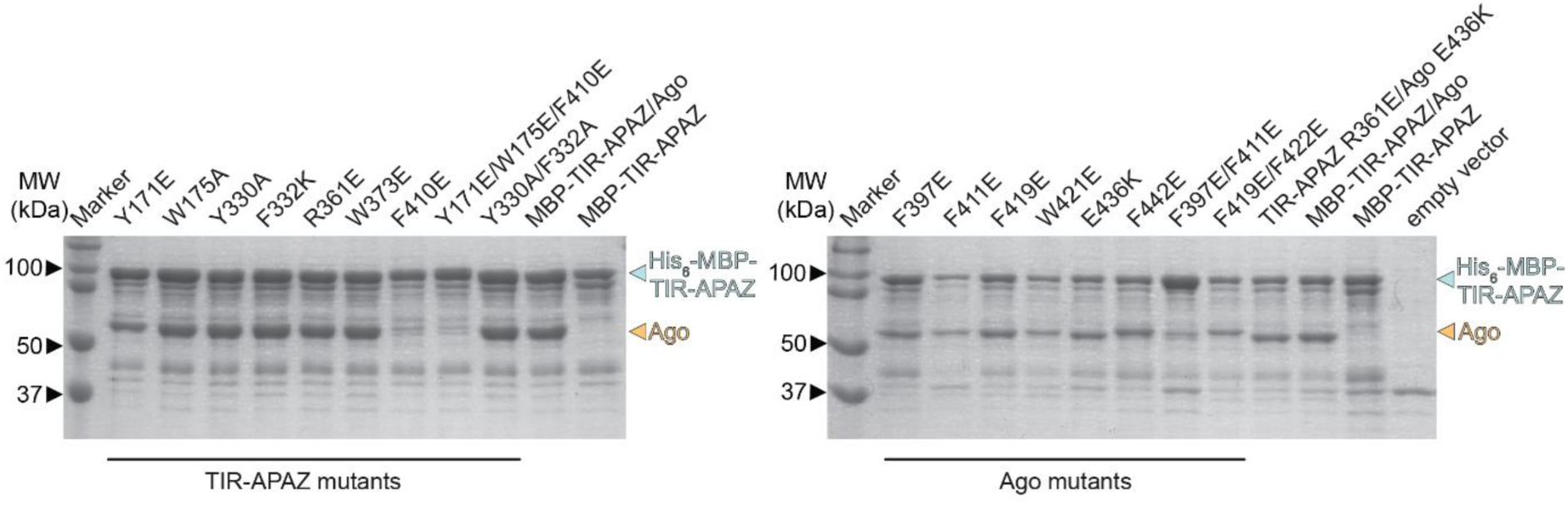
Multiple residues in at the PIWI-APAZ interface contribute to heterodimerization of TIR-APAZ and short pAgo. 6xHis-MBP-BabTIR-APAZ or mutants thereof (left panel), were co-expressed in *E. coli* with BabAgo or mutants thereof (right panel) and proteins were purified using amylose affinity chromatography. Samples were resolved on SDS-PAGE gels, which were used to determine the short BabAgo/BabTIR-APAZ ratio (shown in Figure 1).

**Supplementary Figure S4.**
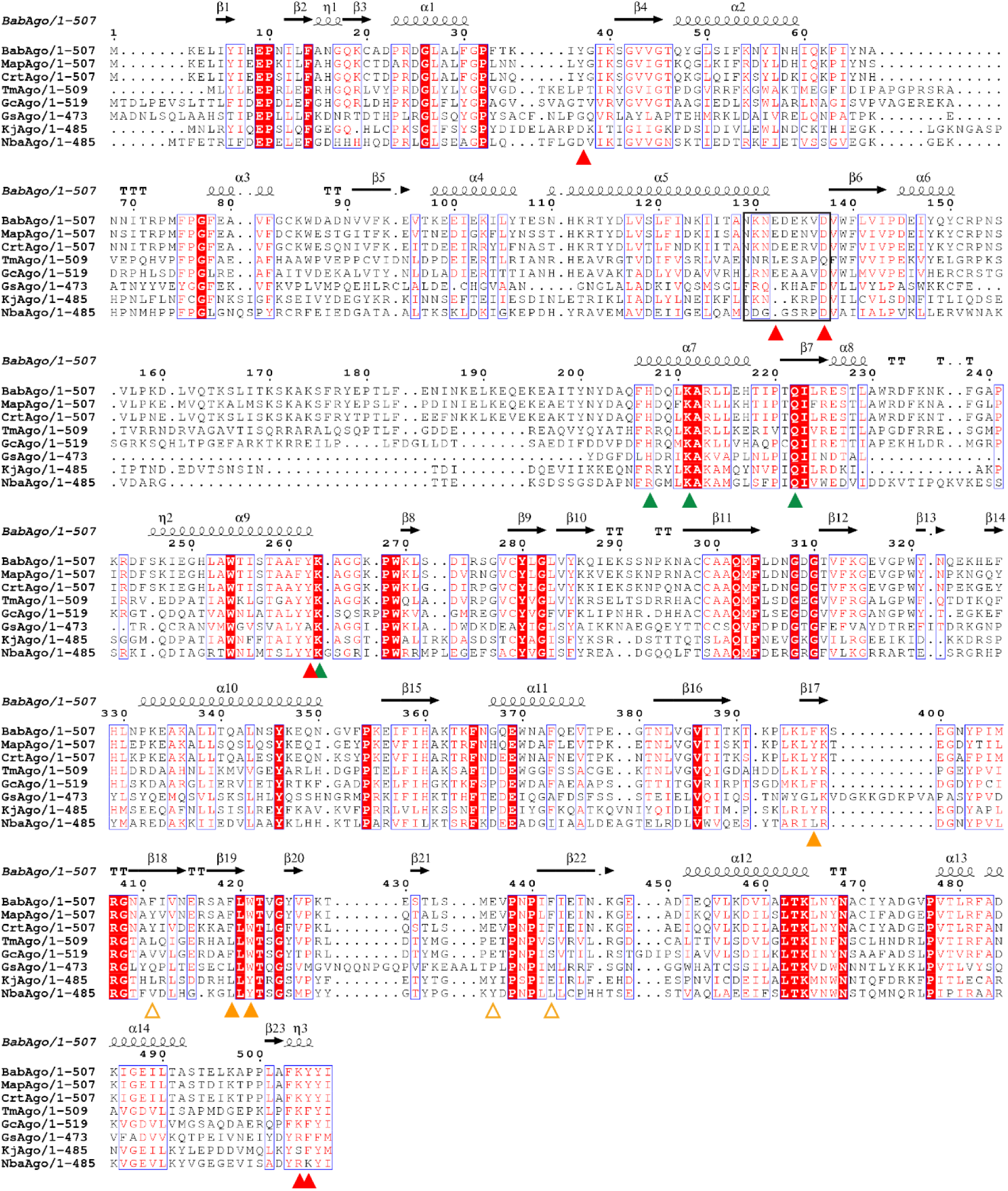
Multiple sequence alignment of short pAgos. Bab: *Bacillales bacterium*; Map: *Maribacter polysiphoniae*^2^; Crt: *Crenotalea thermophila*^2^; Tm: *Tistrella mobilis*; Gc: *Gemmobacter caeruleus*; Gs: *Geobacter sulfurreducens*^3^; Xav: *Xanthomonas vesicatoria*^2^; Kj: *Kordia jejudonensis*^4^; Nba: *Novosphingopyxis baekryungensis*^5^. ▲: Ago-Ago interface residues important for NADase activity. ▲: Ago-APAZ interface residues important for heterodimer complex formation. △: Ago-APAZ interface residues not essential for heterodimer complex formation. ▲: Conserved guide 5′-end phosphate binding residues.

**Supplementary Figure S5.**
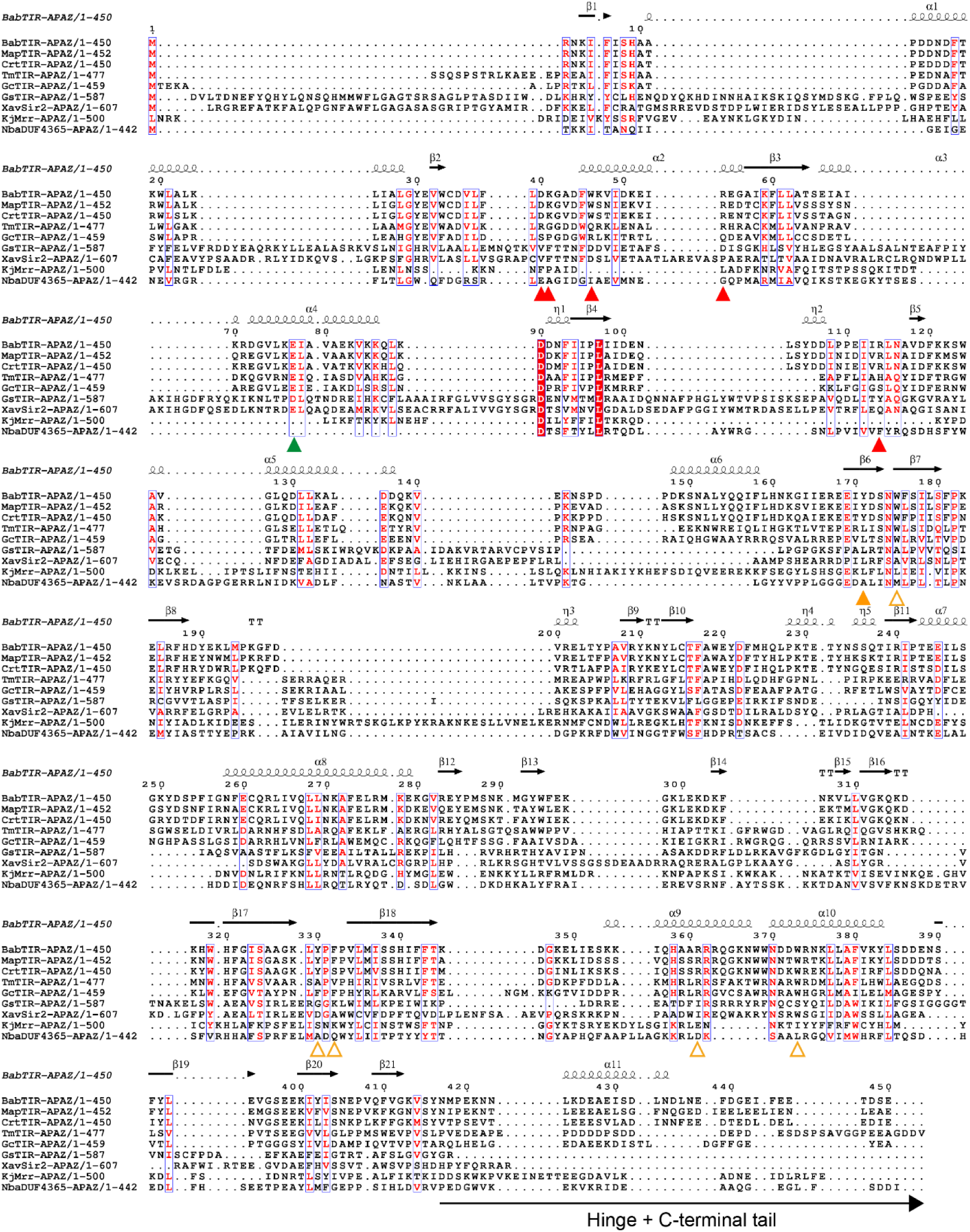
Multiple sequence alignment of APAZ-domain containing proteins. Bab: *Bacillales bacterium*; Map: *Maribacter polysiphoniae*^2^; Crt: *Crenotalea thermophila*^2^; Tm: *Tistrella mobilis*; Gc: *Gemmobacter caeruleus*; Gs: *Geobacter sulfurreducens*^3^; Xav: *Xanthomonas vesicatoria*^2^; Kj: *Kordia jejudonensis*^4^; Nba: *Novosphingopyxis baekryungensis*^5^. Please note that GsSir2-APAZ, XavSir2-APAZ, KjMrr-APAZ, and NbaDUF4365-APAZ have distinct N-terminal effector domains as the other (TIR-APAZ) proteins, and that therefore the first part of alignment should be ignored for these proteins. The APAZ domain starts at BapTIR-APAZ residue 166. ▲: TIR-TIR interface residues important for NADase activity. ▲: Ago-APAZ interface residues important for heterodimer complex formation. Δ: Ago-APAZ interface residues not essential for heterodimer complex formation. ▲: Catalytic residue mutated in catalytic mutants.

**Supplementary Figure S6.**
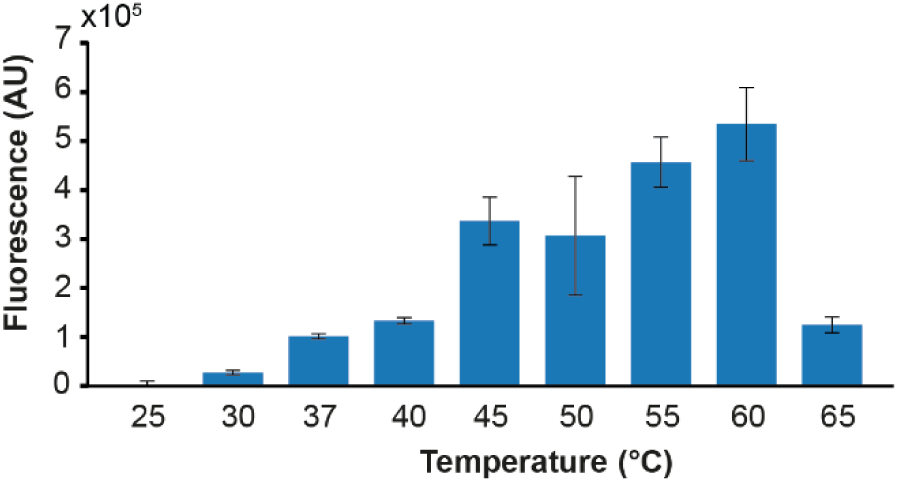
The NADase activity of BabSPARTA at different temperatures. Purified BabSPARTA was mixed with guide RNA and complementary target ssDNA and incubated with ε-NAD+ at different temperatures. After one hour, the fluorescence was measured. The averages of three biological replicates are shown, error bars in bar charts and shadings in growth curves indicate standard deviations. *p < 0.05, **p < 0.01; ***p < 0.001.

**Supplementary Figure S7.**
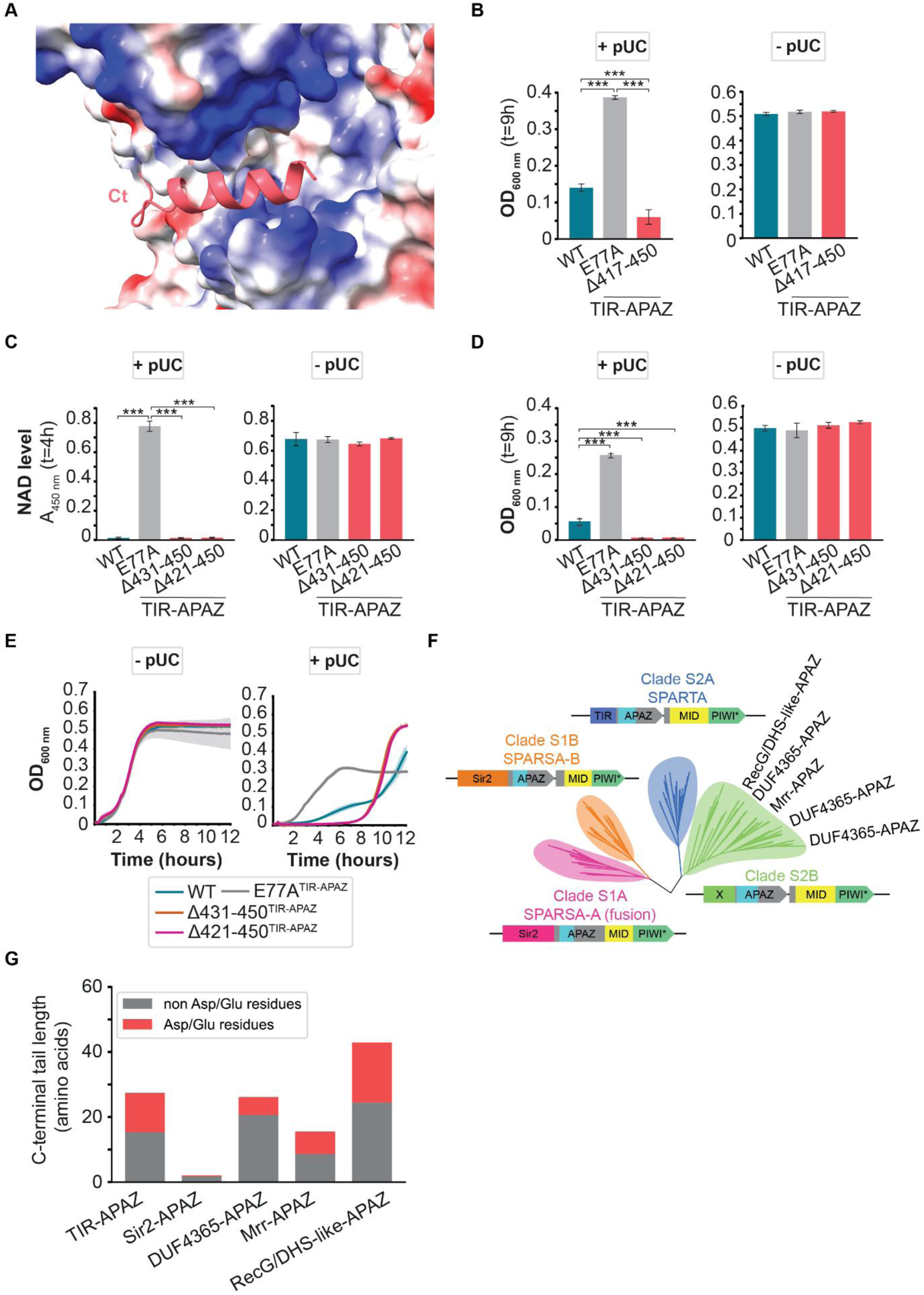
C-terminal tail truncations result in hyperactive BabSPARTA. (**A**) Zoomed-in view of the C-terminal tail region of BabSPARTA. The C-terminal tail is displayed in cartoon representation (in red); the rest of protein is depicted as surface representation colored according to its Coulombic electrostatic potential calculated with ChimeraX (default coloring ranging from red for negative potential through white to blue for positive potential). (**B-E**) Effect of C-terminal tail truncations on SPARTA activity. In the presence (B-D: left panels, E: right panel) or absence (B-D: right panels, E: left panel) of a highly transcribed high copy number plasmid (pUC-mRFP^ΔRBS^), the total NAD (NAD^+^ + NADH) level (C) and OD600 nm (B, D, E) were measured in *E. coli* cultures expressing MapSPARTA, catalytic mutant MapSPARTA^TIR-E77A^, or C-terminal tail truncation mutants Δ417-450^TIR-APAZ^ (B), Δ421-450^TIR-APAZ^ or Δ431-450^TIR-APAZ^ (C-E). Panels B, C, and D show the OD600 nm determined 9 hours after induction of MapSPARTA expression, panel E shows the OD600 nm measured over time starting at 4 hours after induction of MapSPARTA expression. The averages of three technical replicates are shown, error bars indicate standard deviations. (**F**) Schematic unrooted phylogenetic tree containing short pAgos (based on ^6^). (**G**) Conservational analysis of C-terminal extensions of APAZ domain-containing proteins from distinct short pAgo clades. Each bar represent the average C-terminal tail length and average count of Asp/Glu residues for APAZ domain-containing proteins from this clade.

**Supplementary Figure S8.**
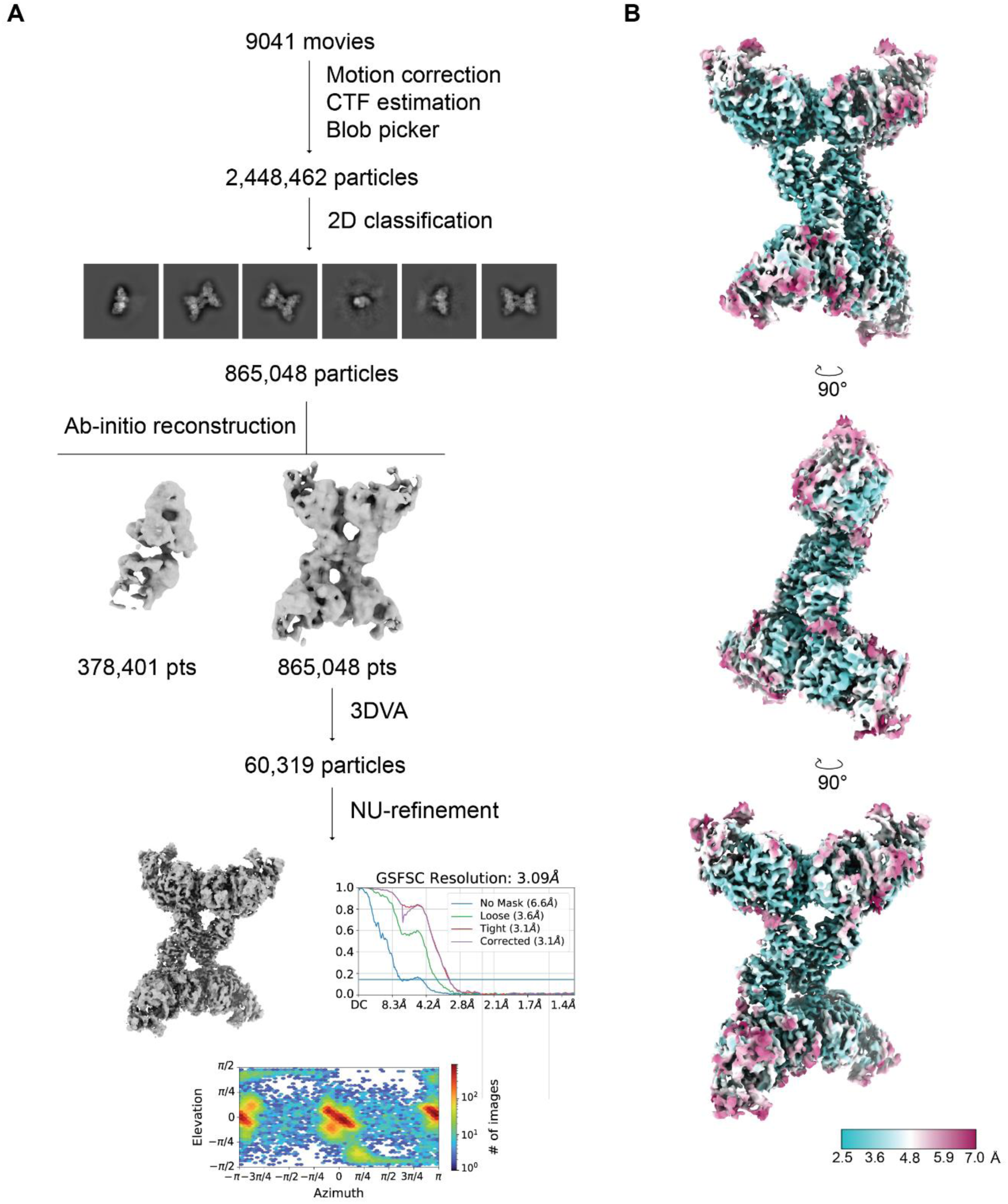
Cryo-EM data processing for BabAgo/BabTIR-APAZ:RNA:DNA complex. (**A**) Cryo-EM image processing workflow for BabAgo/BabTIR-APAZ:RNA:DNA. (**B**) Cryo-EM densities of the BabAgo/BabTIR-APAZ:RNA:DNA complex colored according to local resolution.

**Supplementary Figure S9.**
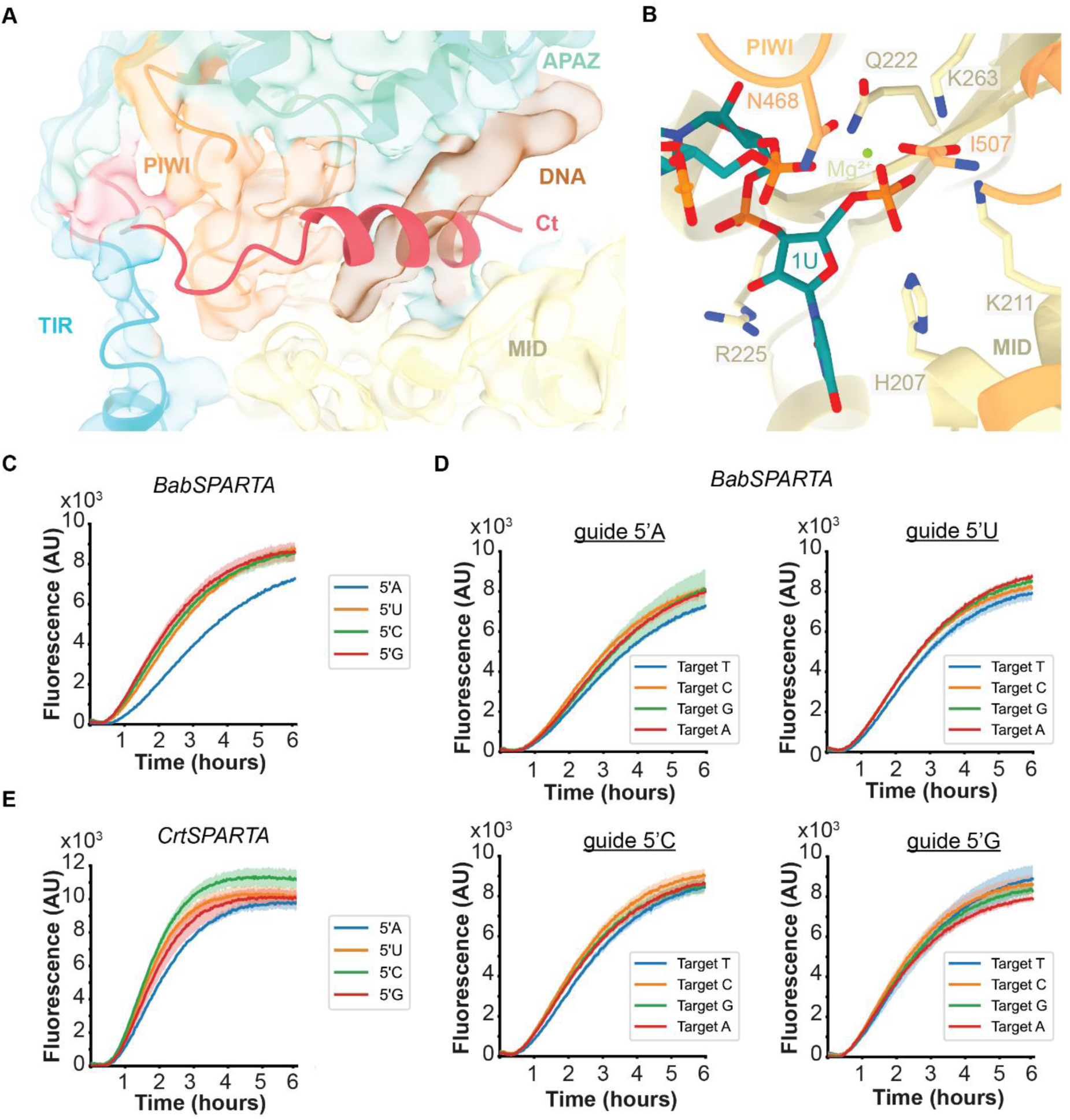
BabSPARTA RNA/DNA duplex coordination. (**A**) Ribbon model of BabSPARTA structure fitted in BabSPARTA:RNA:DNA cryo-EM density, showing that the target bound state lacks any density for the C-terminal tail helix. Density for the RNA/DNA duplex is present in the same region instead. (**B**) Coordination of 5′ end phosphate group of the guide RNA (in teal) by a magnesium ion (in lime green) and residues in the PIWI (in orange) and MID (in yellow) domains. (**C-D**) BabSPARTA can be activated by guide RNAs with all possible 5′ end nucleotides and fully complementary DNA targets (C) as well as DNA targets that have nucleotide substitutions in the position opposite of the 5′ end nucleotide of the guide (D). (**E**) Also CrtSPARTA can be activated by guide RNAs with all possible 5′ end nucleotides. All measurements are corrected with a control without target ssDNA. The averages of three technical replicates are shown, shadings indicate standard deviations.

**Supplementary Figure S10.**
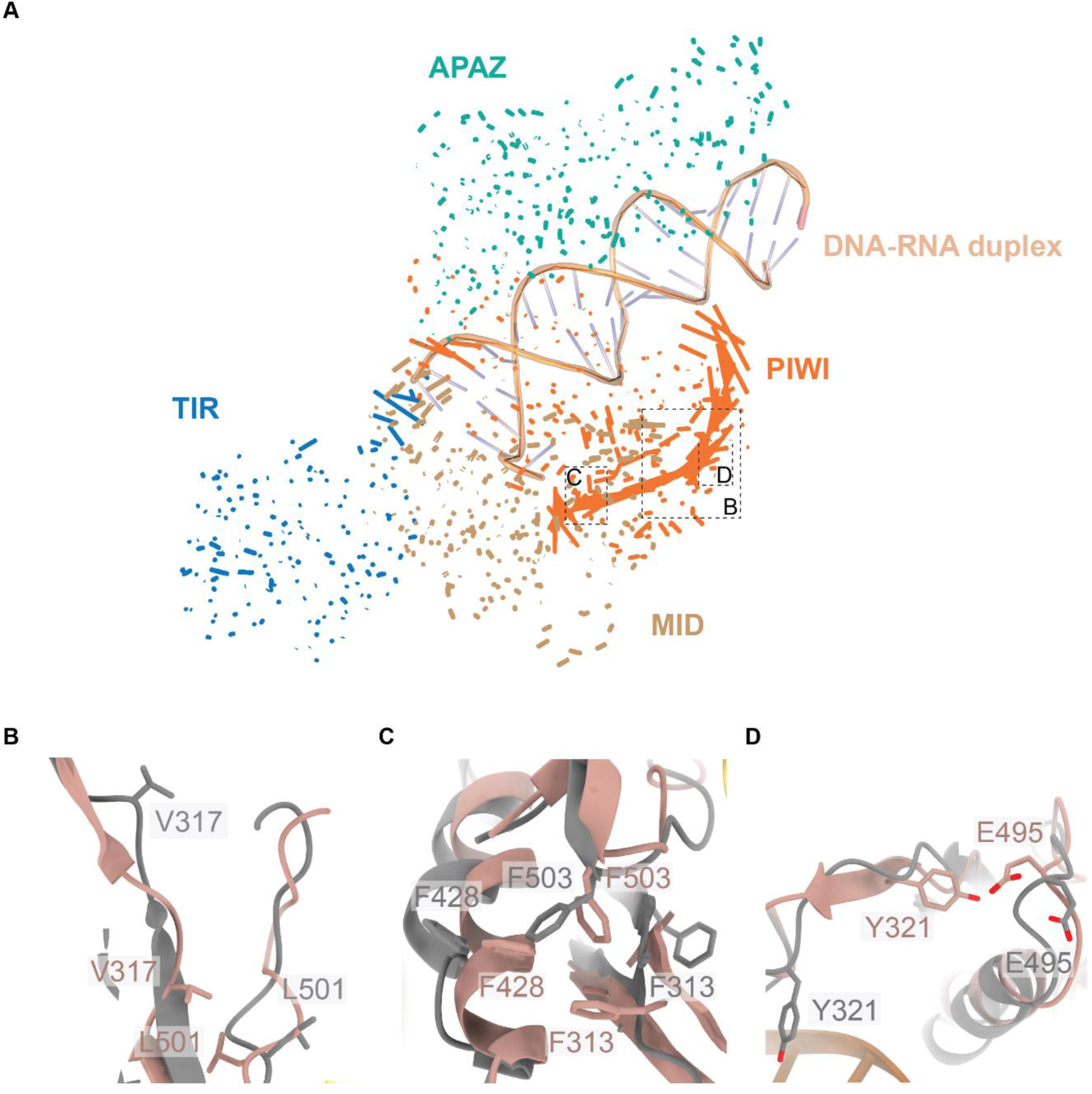
Target binding elicits movements in the PIWI domain that aid dimer formation. (A) The direction of motion between apo and one of the asymmetric copy of target-bound BabSPARTA:RNA:DNA is displayed with vectors of different length. The magnitude of the vectors directly correlates with the amount of motion in that region. Image generated using the modevectors function of PyMOL^1^. (B-D) Rearrangements of PIWI domain residues between apo state (in gray) and tetrameric complex (in pink).

**Supplementary Figure S11.**
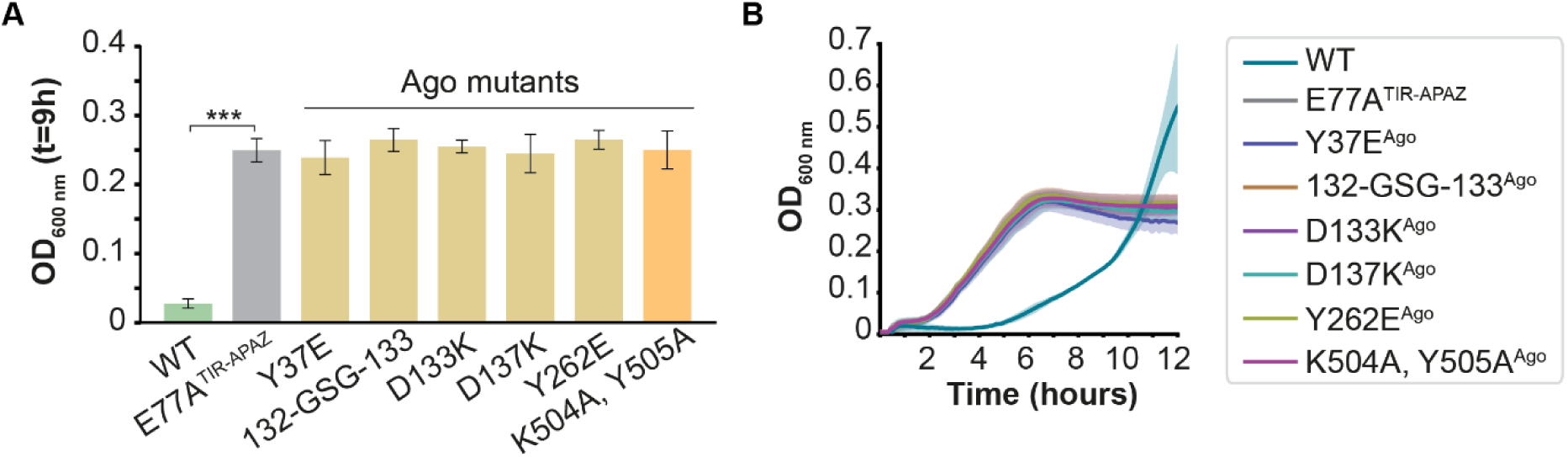
Mutational analyses of SPARTA residues important for catalytic activation. (**A-B**) pAgo-pAgo interaction residues are crucial for catalytic activation of SPARTA. OD600 nm was measured 9 hours after induction of MapSPARTA expression (A) or over time (B) in *E. coli* cultures expressing MapSPARTA, catalytic mutant E77A^TIR-APAZ^, or MapSPARTA with mutated residues at the Ago-Ago interface, in the presence of a highly transcribed high copy number plasmid (pUC-mRFP^ΔRBS^). The averages of three biological replicates are shown, error bars in bar charts and shadings in growth curves indicate standard deviations. *p < 0.05, **p < 0.01; ***p < 0.001.

**Supplementary Figure S12.**
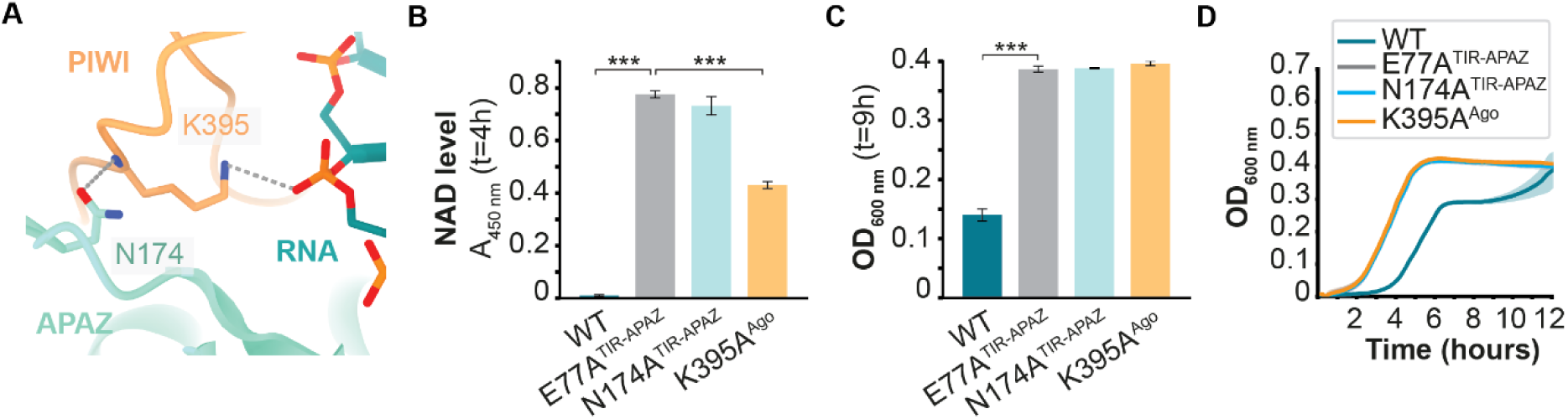
Conserved residues involved in RNA backbone coordination are important for NADase activity of BabSPARTA. (**A**) Zoomed-in view of the interaction between Asn174^APAZ^, Arg395^PIWI^ and the nucleotide-6 phosphate group of the guide RNA. (**B-D**) Interaction of Ago^K395^ with the guide RNA and TIR-APAZ^N174^ is essential for SPARTA activation. The total NAD (NAD^+^ + NADH) level (B) and OD600 nm (C-D) were determined in *E. coli* cultures expressing MapSPARTA, catalytic mutant MapSPARTA^TIR-E77A^, or MapSPARTA^Ago-K395A^, in the presence of a highly transcribed high copy number plasmid (pUC-mRFP^ΔRBS^). Panel C shows the OD600 nm determined 9 hours after induction of MapSPARTA expression, panel D shows the OD600 nm over time starting at 4 hours after induction of MapSPARTA expression. The MapSPARTA and catalytic mutant E77A^TIR-APAZ^ controls are identical to those as in Figure 2 panels B and C and Figure S7 panel B (same experiments). The averages of three biological replicates are shown, error bars in bar charts and shadings in growth curves indicate standard deviations. *p < 0.05, **p < 0.01; ***p < 0.001.

**Supplementary Figure S13.**
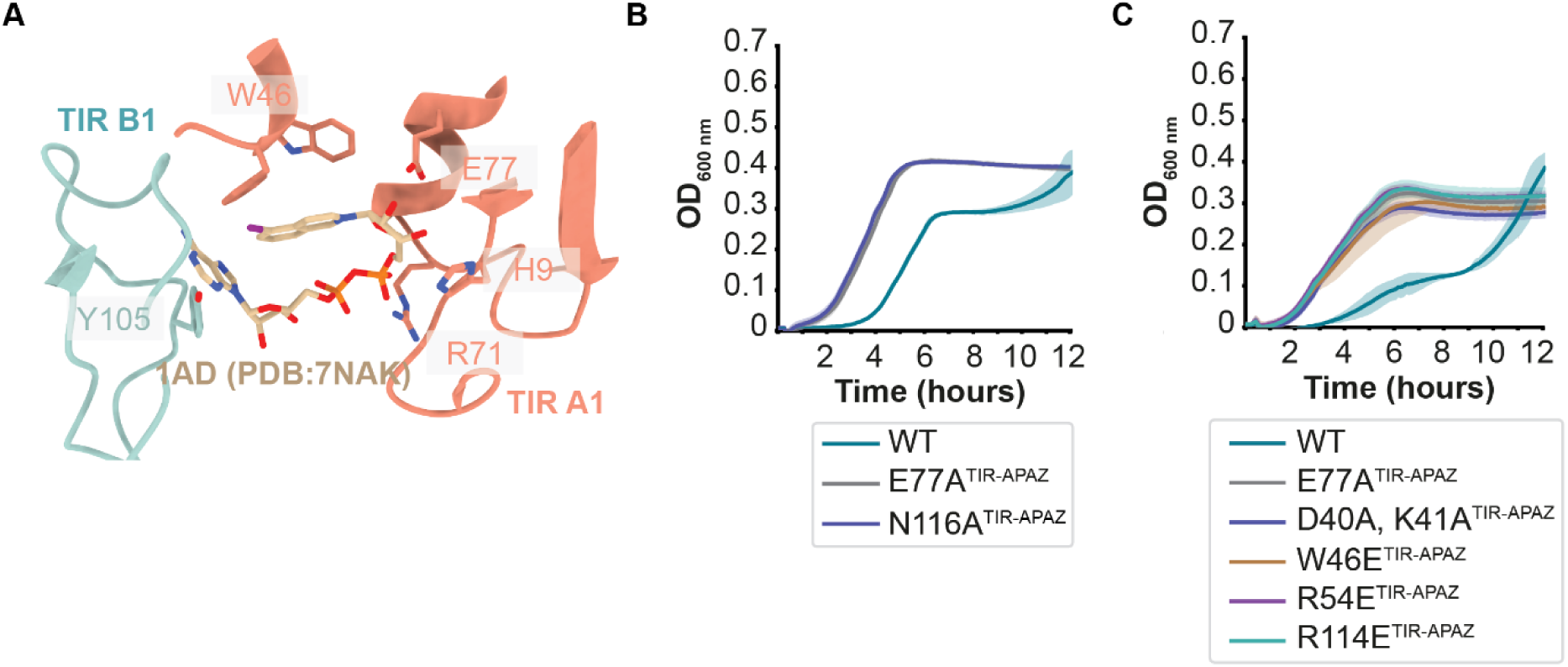
TIR-TIR interactions are essential for BabSPARTA activity. (**A**) Alignment of BabSPARTA (in ribbon representation) with hSARM1 (not displayed) bound to 1AD (in beige, displayed as atom representation) reveals that the ligand is accommodated within a cavity enriched with conserved residues. (**B-C**) TIR-TIR interaction residues are crucial for catalytic activation of SPARTA. OD600 nm was measured or over time in *E. coli* cultures expressing WT MapSPARTA, catalytic mutant E77A^TIR-APAZ^, or MapSPARTA with mutated residues at the TIR-TIR interfaces, in the presence of a highly transcribed high copy number plasmid (pUC-mRFP^ΔRBS^). The MapSPARTA and E77A^TIR-APAZ^ controls that are in the graph together with N116A^TIR-APAZ^ are identical to those in Figure 2 panel C (same experiment). The averages of three biological replicates are shown, error bars in bar charts and shadings in growth curves indicate standard deviations. *p < 0.05, **p < 0.01; ***p < 0.001.

